# Rho family GTPase crosstalk through shared positive and negative feedback mechanisms underlies cytokinetic F-actin waves

**DOI:** 10.64898/2026.11.19.767933

**Authors:** Aamani Thulluru, Jennifer Landino

## Abstract

Cortical excitability, characterized by the rapid polymerization and de-polymerization of cortical F-actin, has been reported in a variety of systems and cell types. In *X. laevis, C. elegans,* and echinoderm embryos, these oscillations are dependent on the small GTPase, Rho, which is regulated in turn by F-actin dependent negative feedback. We sought to investigate whether the Rho family GTPases, Cdc42 and Rac, contribute to specific regulation of F-actin waves and what feedback drives these dynamics. Using an “artificial cortex” system consisting of *Xenopus* egg extract and supported lipid bilayers, we showed that active Cdc42 and active Rac are dynamically patterned in a manner that is coordinated with active Rho and F-actin, suggesting that they contribute to the previously defined oscillatory circuit. We found that total Cdc42 and Rac cortical activity depends on Rho activity. Conversely, Rho activity and F-actin wave dynamics are modulated by Cdc42 and Rac activity. We determined that Cdc42 and Rac are differentially regulated by known regulators of the oscillatory circuit, including Ect2 and F-actin polymerization. We also observed that Rho and Cdc42 are segregated into defined activity zones by microtubule plus-end overlaps, presenting a possible mechanism for the division plane-specific F-actin wave dynamics. Our findings demonstrate that the major Rho family members participate in the cortical excitability circuit and reveal additional mechanisms that regulate Rho and F-actin waves. Additionally, this work supports a model wherein spatial segregation of Rho family GTPases differentially modulates cortical pattern dynamics.

## Introduction

The cell cortex—the actomyosin network directly beneath the plasma membrane of animal cells— mediates cell shape change in a number of essential processes, like cell migration and wound healing (Svitkina, 2018). The Rho family GTPases, Rho, Rac, and Cdc42, promote cortical F-actin assembly by activation of F-actin polymerization factors, like formins and Arp2/3, which drives changes in cell shape (Bement et al., 2006, 2024; Pollard, 2016). Temporally regulated, local activation of Rho GTPases supports the establishment of specific subcellular F-actin structures, enabling a diversity of cellular behaviors (Bement et al., 2006, 2024). In recent years, a phenomenon termed “cortical excitability” has been observed across many systems, and emerges during processes defined by cell shape change (Vicker, 2000; Bement et al., 2015; Michaux et al., 2018). This phenomenon consists of rapid and meso-scale waves of F-actin polymerization, and is generated through positive and negative feedback between F-actin and its effectors (Bement et al., 2015; Michaux et al., 2018; Michaud et al., 2022). In echinoderm and frog embryos, these oscillations are driven primarily by Rho, which recruits its own activator, Ect2 and stimulates F-actin polymerization (Michaud et al., 2022). F-actin in turn recruits the Rho inhibitor RGA3/4, leading to an excitable activator-inhibitor circuit of Rho and F-actin oscillations (Michaud et al., 2022; Figure S1E). It is not known whether Cdc42 and Rac are regulated by similar positive and negative feedback mechanisms and coordinated with these Rho and F-actin waves, nor how Rho and F-actin waves support cell shape change.

F-actin waves are spatially regulated in the developing embryo, with wave dynamics dampened at the division plane compared to other cortical regions (Swider et al., 2022). The mechanism of spatial regulation of F-actin waves is not well understood. One possible explanation is that equatorial enrichment of Rho activity and F-actin assembly (Green et al., 2012) may be sufficient to drive division plane-specific wave dynamics. An alternative explanation is the Rho family GTPases may be spatially patterned in the cell, as has been described for cell migration (Bement et al., 2006; De Belly et al., 2026) and wound healing (Benink and Bement, 2005; Vaughan et al., 2011). Previous investigations of Rho family GTPase wave patterning in other systems has posited that that feedback circuits rapidly coordinate local and global GTPase activity and Rho family GTPase cross talk (Iglesias and Devreotes, 2012; Kuhn et al., 2025; De Belly et al., 2026). For example, Rho and Rac waves in migrating T-cells display long-range feedback *via* the plasma membrane which supports spatially-coordinated activation of Rac and Rho despite their spatial separation (De Belly et al., 2026). It is unknown whether similar feedback between the Rho family GTPases, Rho, Rac, and Cdc42, drives specific cytokinetic F-actin wave dynamics.

Here we investigate the spatiotemporal relationship between Rho family GTPases in excitable patterning as well as the role of Rac and Cdc42 in regulating F-actin waves. We use an *in vitro* “artificial cortex” system made up of *Xenopus* egg extract and a supported lipid bilayer (SLB) which has been previously shown to reconstitute active Rho and F-actin wave patterning (Landino et al., 2021). We find that active Rac and active Cdc42 are spatially and temporally coordinated with dynamic waves of Rho and F-actin in the artificial cortex system. The emergence of Rac and Cdc42 excitable wave patterning suggests that they are regulated by both positive and negative feedback. We find shared mechanisms of positive feedback, mediated by Ect2, and negative feedback, mediated by F-actin, between the Rho family GTPases. Additionally, we find that the Rho family GTPases regulate each other’s activity, with Rho mediating total GTPase activity and Cdc42 and Rac regulating Rho and F-actin wave dynamics. Finally, we discover novel spatial organization of the Rho family GTPases, induced by microtubule plus end overlaps, which are key spatial markers of cytokinesis (Nguyen et al., 2014). This finding provides a possible mechanism to explain division plane-specific F-actin wave dynamics.

## Results

### Cdc42 and Rac are present and active in the artificial cortex

We first sought investigate the relative cortical activity of Rho, Rac, and Cdc42 in our artificial cortex system (Landino et al., 2021). We quantified the total GTPase activity and F-actin assembly on SLBs over time, using Total Internal Reflection Fluorescence (TIRF) microscopy. Biosensors for active Cdc42 and active Rac (Cdc42 binding domain of N-WASP, wGBD; Rac binding domain of Rhotekin, pGBD) were used to measure Cdc42 and Rac activity relative to active Rho and F-actin (Rho binding domain of Rhotekin, rGBD; Utrophin calponin homology domain, UtrCH) (Bement et al., 2006). Notably, previous studies have shown that these biosensors are specific for the membrane-localized active pool of the respective GTPase (Golding et al., 2019; Armstrong et al., 2025). We hereafter use the simplified Rho, Cdc42, or Rac nomenclature to refer to the active form of the GTPase.

We first investigated the initial rate of cortical activation of Cdc42 and Rac relative to Rho and F-actin in the artificial cortex (Figure S1A-C). We added egg extract to SLBs during a live imaging time course and quantified total intensity as a readout of GTPase activation during cortex assembly. We found that Rac and Cdc42 robustly localize to the artificial cortex SLB and activity peaked after maximal Rho and F-actin signal (Figure S1A-C). This suggests that Cdc42 and Rac activation during cortex assembly is downstream of Rho-mediated cortical actin assembly. We next investigated their sustained activity during an hour-long time course. Total Rho intensity stayed relatively constant over this time period (Figure S2A); however, Cdc42 and Rac decrease over time (Figure S2A). Cdc42 signal decreases early in the time course and stays relatively constant thereafter. Rac, however, is particularly short-lived, with half of the starting intensity lost after the first twenty minutes of the reaction, followed by a continued decline (Figure S2A). Within individual experiments, we found that Rac localization on the SLB was temporally anti-correlated with Rho and F-actin (Figure S2C), suggesting that Rac may be negatively regulated by Rho activity in the artificial cortex, as has been observed in cell migration (De Belly et al., 2026).

### Active Cdc42 and active Rac participate in Rho-mediated F-actin traveling waves

In the artificial cortex system, cortical dynamics are organized into two types of patterns: traveling waves and standing oscillations (Landino et al., 2021). Traveling waves move directionally across the SLB, while standing oscillations emerge from periodic cycles of activation and inactivation in the same place. Both types of waves display similar temporal dynamics, with Rho activity preceding F-actin polymerization. We observed both Cdc42 and Rac localized to traveling waves of Rho and F-actin in the artificial cortex (Figure 1A-B, D-E). To quantify the spatiotemporal relationship between Rho family GTPases and F-actin in traveling waves, we measured biosensor intensity over time over a small (4 x 4 μm) region in the field of view as the traveling wave passed through. We found that Cdc42 activity peaks between Rho and F-actin (Figure 1C), suggesting Cdc42 activity is downstream of Rho in traveling waves. Rac displayed an unexpected temporal relationship with Rho and F-actin, with Rac activity preceding Rho (Figure 1F). Furthermore, Rac activity is unusually sustained, with peak activity persisting until after peak Rho activity (Figure 1F). To better understand the relationship between Cdc42, Rac, Rho and F-actin, we performed high-throughput analysis of wave dynamics (Github, *waveanalysis*, Figure S1F-G) to quantify the offset between activity peaks (temporal shift, Figure S1G). We confirmed that all three of the Rho family GTPases act upstream of F-actin in traveling waves (Figure 1G, Figure S2D-E). This analysis also confirmed that Cdc42 acts downstream of Rho, while Rac acts upstream of Rho activity (Figure 1G, Figure S2D-E). These findings demonstrate that the activity of each Rho family GTPase has defined spatiotemporal coordination relative to its family members and to and F-actin within traveling waves.

**Figure 1.**
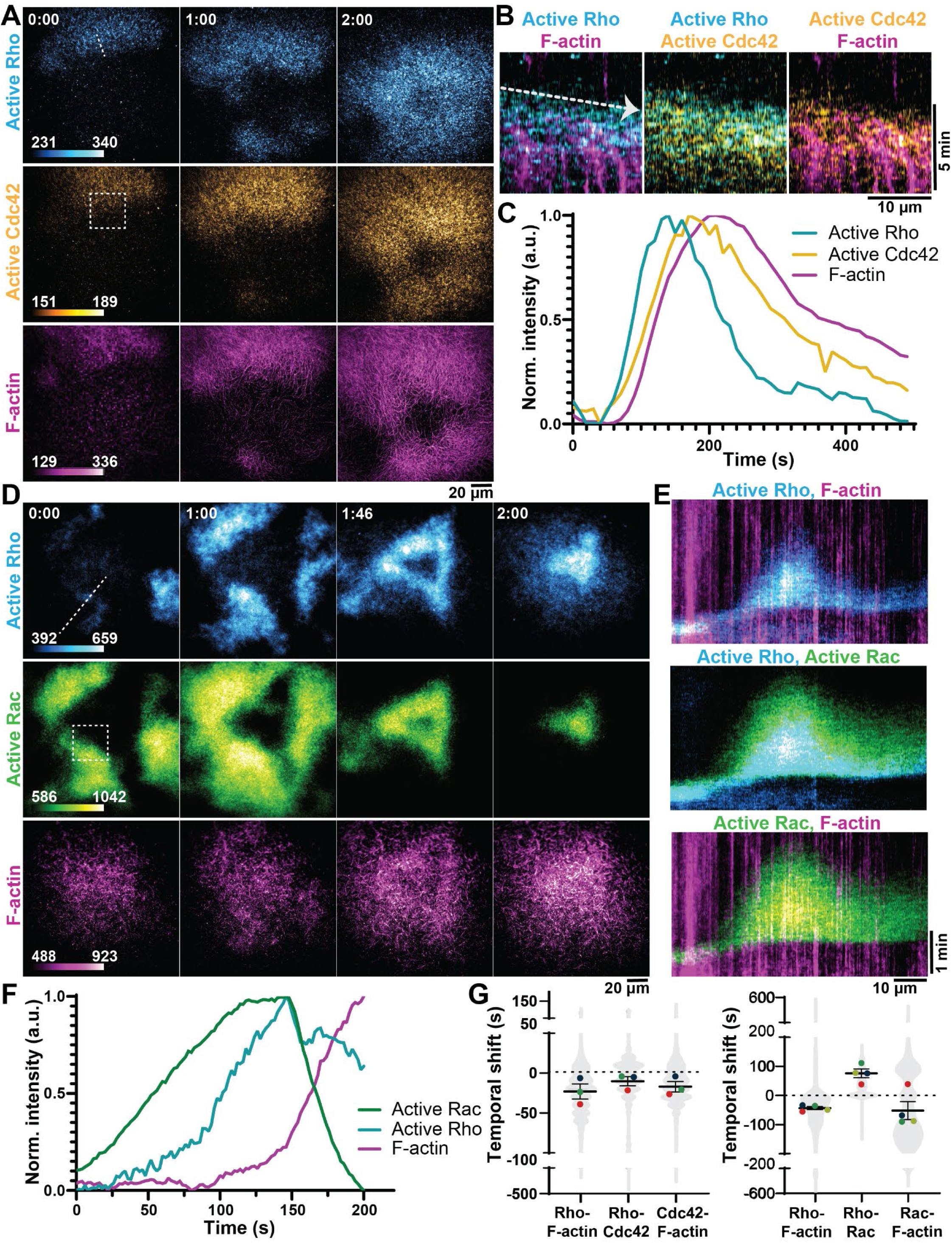
Active Cdc42 and active Rac traveling waves are coordinated with waves of Rho and F-actin. **(A)** Representative micrographs of Rho (cyan), Cdc42 (yellow), and F-actin (magenta) traveling waves over time. Time is indicated in min:sec. Dashed line indicates region used to generate kymographs in **(B)**; box indicates region used for intensity over time quantification in **(C)**. **(B)** Kymographs of Rho and Cdc42 traveling waves. Arrow indicates direction of the wave front. Rho and Cdc42 GTPase signal precedes F-actin polymerization. **(C)** Intensity over time of Rho, Cdc42, and F-actin traveling waves in a 30 x 30 μm region of interest (ROI). **(D)** Representative micrographs of Rho (cyan), Rac (green), and F-actin (magenta) traveling waves over time. Time is indicated in min:sec. Dashed line indicates region used to generate kymographs in **(E)**; box indicates region used for intensity over time quantification in **(F)**. **(E)** Kymographs of Rho and Rac traveling waves. Rho and Rac GTPase signal precedes F-actin polymerization. **(F)** Intensity over time of Rho, Rac, and F-actin traveling waves in a 30 x 30 μm ROI. **(G)** Quantification of the pairwise temporal shift between GTPases and F-actin. Negative shift indicates preceding order. Mean ± standard error (SE) is shown; violins represent all ROIs analyzed for all movies and dots indicate means of individual experiments. Within each graph, dot color indicates replicate. For Cdc42, n = 3 movies, 3 extract preps from 3 frogs, 2350 ROIs. For Rac, n = 3 movies, 3 extract preps from 3 frogs, 3132 ROIs.

### Active Cdc42 and active Rac dynamically oscillate in coordination with Rho and F-actin

We next investigated standing oscillations in the artificial cortex, and found that both Cdc42 and Rac formed oscillatory patterns in a manner that is coordinated with Rho and F-actin standing oscillations (Figure 2A-F). As Rho oscillations have been previously characterized *in vivo* and *in vitro* (period of 50 to 130s; (Bement et al., 2015; Landino et al., 2021; Schwarz et al., 2026), we co-imaged Rho dynamics as a control (period of 60 ± 15 s, Figure 2G-H). Using our custom analysis code, we found that Cdc42 and Rac oscillate with approximately a one-minute period (Figure 2G-H; Cdc42: 60 ± 13 s, Rac: 59 ± 32 s), like Rho. The similar period observed for the Rho family GTPases suggests coordinated positive and negative feedback mechanisms between the family members.

**Figure 2.**
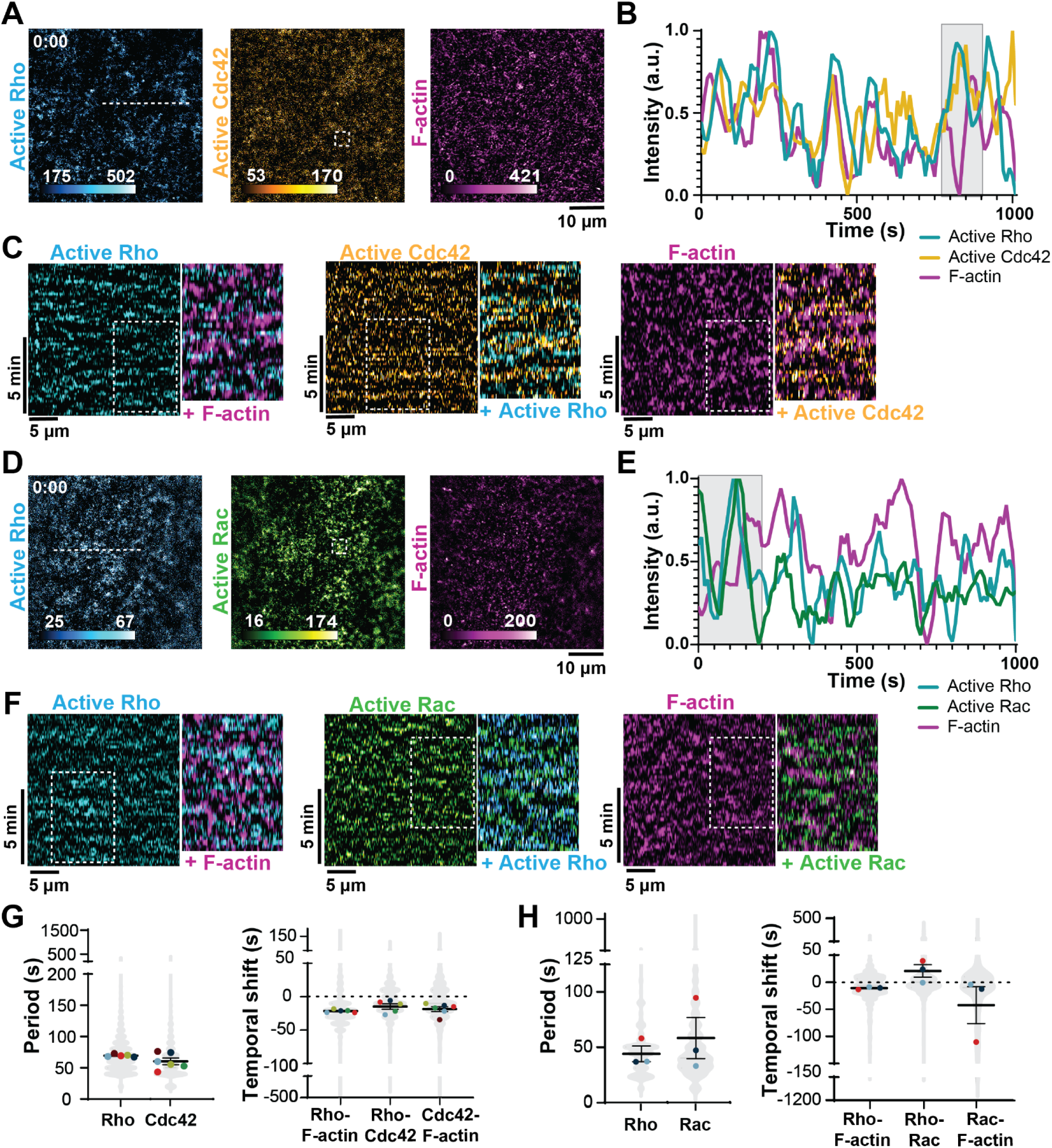
Active Cdc42 and active Rac participate in cortical oscillations in the artificial cortex. **(A)** Representative micrographs of Rho (cyan), Cdc42 (yellow), and F-actin (magenta) oscillations. Dashed line indicates region used to generate kymographs in **(C)**; box indicates region used for intensity over time measurements in **(B). (B)** Representative intensity over time plots showing an example oscillation of Rho, Cdc42, and F-actin (highlighted in gray). **(C)** Kymographs of Rho, Cdc42, and F-actin oscillations showing periodic GTPase signal. Rho and Cdc42 precede F-actin; Rho precedes Cdc42. **(D)** Representative micrographs of Rho (cyan), Rac (green), and F-actin (magenta) oscillations. Dashed line indicates region used to generate kymographs in **(F)**; box indicates region used for intensity over time measurements in **(E)**. **(E)** Representative intensity over time plots showing an example oscillation of Rho, Rac, and F-actin (highlighted in gray). **(F)** Kymographs of Rho, Rac, and F-actin oscillations, showing periodic GTPase signal. Rho and Rac precede F-actin; Rac precedes Rho. **(G)** Quantification of Rho and Cdc42 period and pairwise temporal shifts of Rho, Cdc42, and F-actin. Mean ± SE is shown; violins represent all ROIs analyzed for all movies and dots indicate means of individual experiments. Dot color indicates replicates. n = 6 movies, 6 extract preps from 6 frogs, 4698 ROIs. **(H)** Quantification of Rho and Rac period and pairwise temporal shifts of Rho, Rac, and F-actin. Mean ± SE is shown; violins represent all ROIs analyzed for all movies and dots indicate means of individual experiments. Dot color indicates replicates. n = 3 movies, 3 extract preps from 3 frogs, 2350 ROIs.

We also analyzed the temporal shift between each GTPase and F-actin in standing oscillations. Our control analysis of the Rho/F-actin temporal shift in standing oscillations agreed with previous reports (Landino et al., 2021; Schwarz et al., 2026). Like we observed in traveling waves, we found that Cdc42 and Rac peaked before F-actin, with Cdc42 peaking 19 ± 9 s prior to F-actin peaks (Figure 2G), and Rac peaking 42 ± 59 s prior to F-actin (Figure 2H). Furthermore, we examined the temporal relationship between GTPases and found that the relative temporal dynamics of standing oscillations mirrored our observations from traveling waves. We found that Rho precedes Cdc42 in standing oscillations, with Rho peaking 15 ± 9 s prior to Cdc42 peaks (Figure 2G, Figure S3A), and that Rac precedes Rho, peaking 21.11 ± 20.27 s before Rho (Figure 2H, Figure S3B). Additional wave dynamics of standing oscillations, including peak relative amplitude and peak width, were similar for Rho, Cdc42, and Rac (Figure S3C-D).

Together with our findings in traveling waves, this result demonstrates that Cdc42 and Rac, like Rho and F-actin, are dynamically patterned within the artificial. The defined temporal order of GTPase activation within traveling waves and standing oscillations suggests specific regulatory mechanisms exist to control the relative activity of Rho family GTPases during cortical patterning. Moreover, this oscillatory behavior of Cdc42 and Rac indicates they are regulated by a coupled activator-inhibitor circuit, as previously described for Rho (Bement et al., 2015; Michaud et al., 2022).

### F-actin negatively regulates Rho and Rac, but not Cdc42, activity

As Rho, Rac, and Cdc42 wave dynamics are tightly regulated in space and time, we wanted to investigate whether shared feedback mechanisms could explain coordination of Rho family GTPase activity. Rac and Cdc42 oscillate on a similar timescale to Rho and precede F-actin polymerization. Therefore, we hypothesized that Rac and Cdc42 may be inhibited by F-actin polymerization, as has been shown for Rho (Bement et al., 2015). To test the effects of F-actin on activity of each of the Rho family GTPase, we inhibited F-actin assembly with Latrunculin A, which blocks polymerization through sequestration of actin monomer.

We incubated egg extract with 15 μM LatA prior to addition to the SLB (Figure 3A) and assayed total GTPase activity over time (Figure 3B, D). As expected, we see that F-actin intensity is decreased by LatA treatment (Figure 3B-E) compared to the DMSO vehicle control. We observed an increase in total Rho activity relative to vehicle controls with LatA treatment, confirming that F-actin inhibits Rho activity in the artificial cortex (Figure B-E), but observed only marginal changes to Rho period or amplitude (Figure S4B-C). We also assayed Cdc42 and Rac activity after LatA treatment and found Cdc42 activity decreased while Rac activity increased (Figure 3B-E). Increased Rac activity after F-actin inhibition suggests that Rac, like Rho, is negatively regulated by F-actin in our system. Our finding that Cdc42 activity decreases in response to F-actin inhibition suggests that feedback signaling between Cdc42 and F-actin is more complex. Notably, the observed differences in Cdc42 and Rac activity after F-actin inhibition suggest the regulatory mechanisms are specific to each Rho family GTPase.

**Figure 3.**
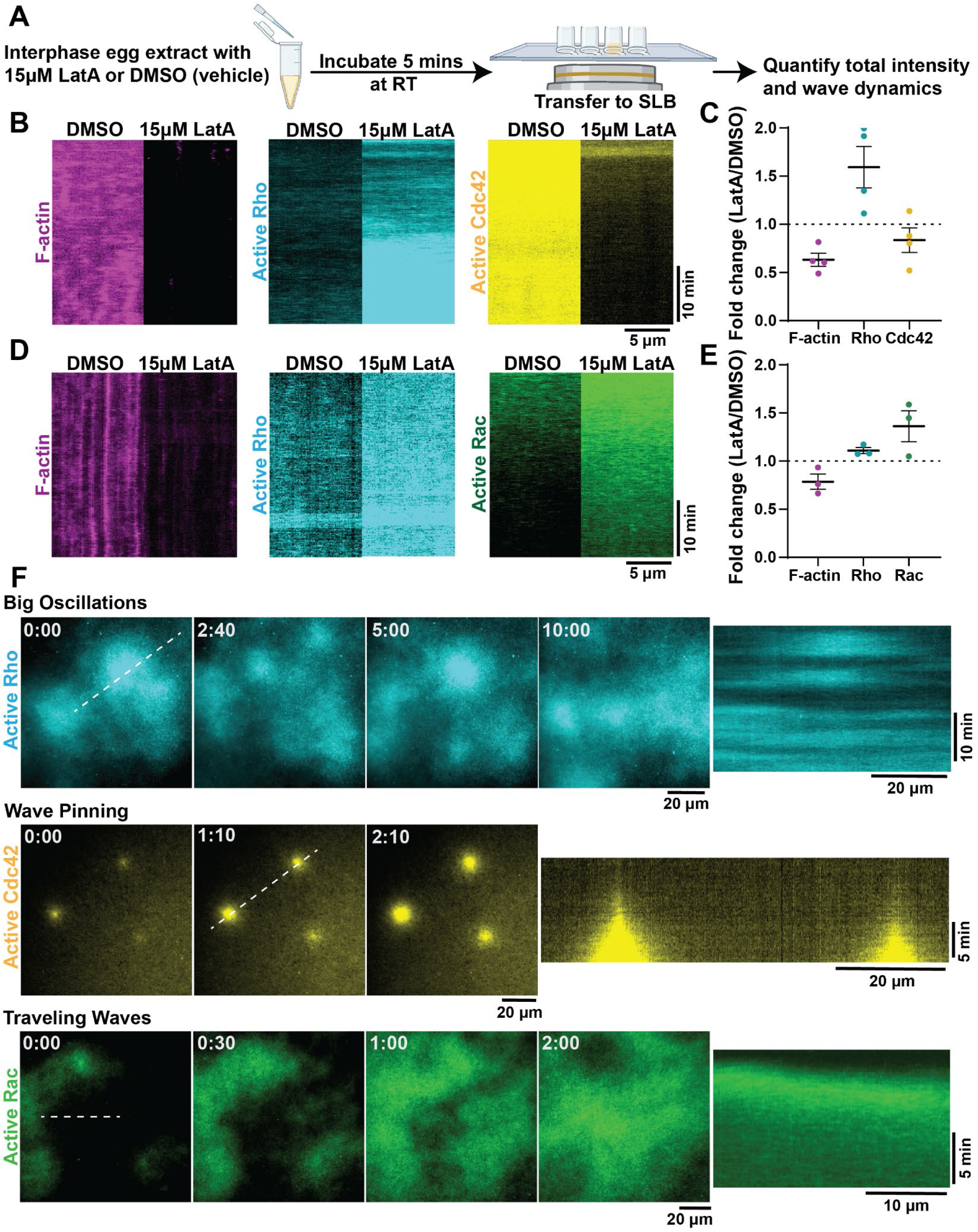
F-actin inhibition activates Rho and Rac GTPases and decreases Cdc42 GTPase activity. **(A)** Schematic of experimental design. 15 µM LatA or DMSO (vehicle control) were added to extract containing fluorescent probes and incubated at room temperature for 5 minutes prior to addition to SLB. **(B)** Representative kymographs of total Rho (cyan), Cdc42 (yellow), or F-actin (magenta) signal after DMSO (left) or 15 μM LatA (right) treatment. Intensity is scale-matched across experimental conditions. **(C)** Average fold change in GTPase and F-actin intensity following DMSO or LatA treatment; dots represent one replicate. Fold change lower than one indicates a decrease in signal intensity following LatA treatment. Mean ± SE is shown. n = 8 movies, 4 extract preps from 4 frogs, 8 ROIs. **(D)** Representative kymographs of Rho (cyan), Rac (green), or F-actin (magenta) after DMSO (left) or 15 μM LatA (right) treatment. Intensity is scale-matched across experimental conditions. **(E)** Average fold change in GTPase and F-actin intensity following DMSO or LatA treatment; dots represent one replicate. Fold change lower than one indicates a decrease in signal intensity following LatA treatment. Mean ± SE is shown. n = 6 movies, 3 extract preps from 3 frogs, 6 ROIs. **(F)** *Left:* Representative micrographs of infrequent wave patterning regimes observed after LatA treatment including big oscillations, wave pinning and traveling waves. Time is indicated in min:sec from the start of the emergence of new patterning. Dotted line indicates regions used to generate kymographs. *Right:* Kymographs of big oscillations, wave pinning, and traveling waves. All Rho family GTPases were observed to localize to these LatA-induced wave patterns. Infrequent wave patterning regimes were observed following LatA treatment in 6/7 experimental replicates, and in 3/7 replicates following DMSO treatment.

We examined the temporal dynamics of Cdc42 and Rac oscillations after LatA treatment and, like Rho, observed only modest differences in the period, amplitude, or temporal shift between GTPase activity peaks compared to vehicle controls (Figure S4B-D). Interestingly, we did observe that oscillatory dynamics in LatA-treated samples were more likely to develop patterning regimes infrequently observed in our system, including traveling waves, wave pinning, and big oscillations (Figure 3F). Big oscillations are characterized by periodic patches of wave activity much larger than what we observe in standing oscillations, spanning tens of microns. We also observed patterns characterized by a gradual radial expansion of GTPase signal from defined foci in a manner that could be analogous to wave pinning (Mori et al., 2008). We observed the emergence of these wave patterns in 6 out of 7 samples that were treated with LatA, and only in 3 out of 7 for DMSO (vehicle) treatment. Although the mechanisms that generate these infrequent wave patterns are not clearly understood, this observation suggests that feedback from F-actin plays a significant role in regulating the excitable patterning regime of Rho family GTPases.

### Ect2 provides positive feedback to Rho and Cdc42, but not to Rac

As excitable Rho and F-actin patterning requires coupled positive and negative feedback, we next sought to determine mechanisms of positive feedback to Cdc42 and Rac. The RhoGEF Ect2 makes up the positive feedback arm of the known cortical excitability circuit, and is recruited and activated by Rho (Chen et al., 2020; Michaud et al., 2022; Figure S1E). Ect2 GEF activity is regulated, in part, by autoinhibition of its DH (GEF) domain by the N-terminal BRCT domains (Chen et al., 2020; Figure 4B). To assay the role of positive feedback signaling in our system, we purified a C-terminal fragment of Ect2 (Ect2-DH), containing its DH (GEF) and PH (membrane-binding) domains (Figure 4B). We predicted that this C-terminal fragment will be constitutively active, as Ect2 autoinhibition requires its N-terminus (Chen et al., 2020).

**Figure 4.**
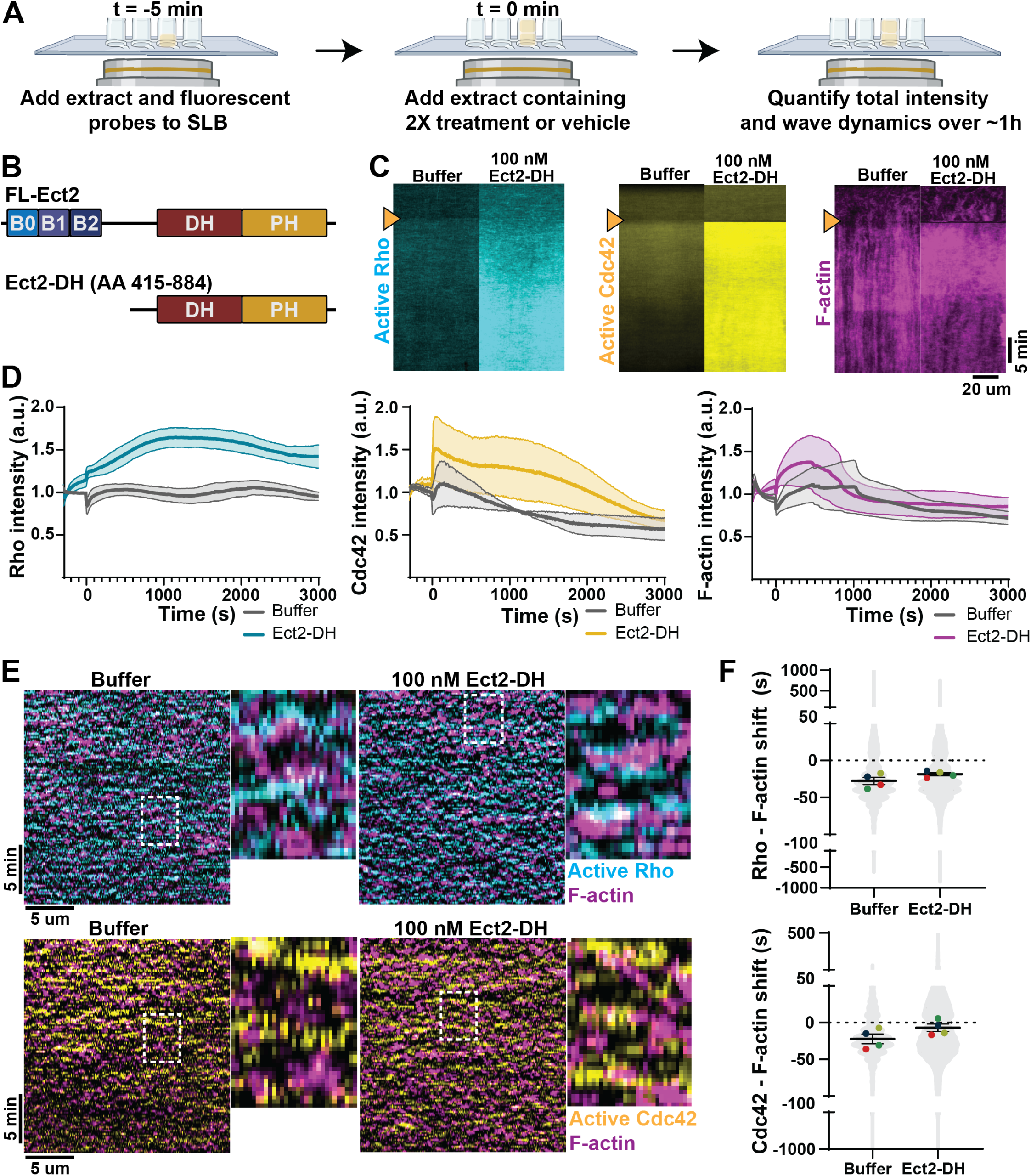
Ect2-DH increases Rho and Cdc42 activity and alters wave dynamics. **(A)** Schematic of experimental setup. Untreated extract containing fluorescent probes was imaged for 5-10 min prior to addition of treatment or vehicle control. **(B)** Schematic of full length Ect2 and Ect2-DH truncation used to generate constitutively active Ect2. **(C)** Representative kymographs of Rho, Cdc42, or F-actin intensity over time. Orange arrowheads indicate time of buffer or Ect2 addition. Intensity is scale-matched across experimental conditions. **(D)** Quantification of mean intensity over time. Solid line represents mean, shaded area represents SE. n = 6 movies, 3 extract preps from 3 frogs, 6 ROIs. T = 0 is time of buffer or Ect2-DH addition. **(E)** Representative kymographs of cortical oscillations following treatment with Ect2. Zoomed regions (right) are highlighted by white boxes. GTPase peaks and F-actin peaks become closer together following Ect2-DH addition. **(F)** Quantification of temporal shift between GTPase activity and F-actin upon buffer or Ect2-DH addition. Mean ± SE is shown; violin represents all regions of interest analyzed in all movies, dots indicate means of individual experiments. Dot color indicates replicates. n = 6 movies, 3 extract preps from 3 frogs, 4700 ROIs.

We first conducted a gelation-contraction assay of egg extract to evaluate Ect2-DH activity in our system (Field et al., 2011). Using droplets of egg extract in oil, we confirmed that interphase egg extract treated with a buffer control was not contractile, as expected (Figure S5A). Addition of 250 nM Ect2-DH was sufficient to induce rapid contraction of interphase extract, as evidenced by the condensed cytoplasmic mass within the droplet (Figure S5A). This result indicates that the truncated Ect2-DH fragment is sufficient to drive increased Rho-mediated actomyosin contractility in egg extract, suggesting Ect2-DH is hyperactive (Jaffe and Hall, 2005).

To investigate the role of Ect2-dependent positive feedback in regulating Rho family GTPases, we assayed GTPase activity in response to Ect2DH addition in the artificial cortex system. We used a lower concentration of Ect2-DH (100 nM) to prevent bulk contraction of the cytoplasm. We first allowed the cortex to assembly, then added Ect2-DH during the time course, and assayed GTPase activity over time (Figure 4A). We found that addition of Ect2-DH caused both Rho and Cdc42 activity to increase (Figure 4C-D), while Rac activity and F-actin polymerization were unchanged (Figure S5B, Figure 4C-D). This finding agrees with previous work showing that Ect2 activates cortical Rho activity (Bement et al., 2015), and demonstrates that Ect2 hyperactivation also positively regulates Cdc42.

We analyzed oscillation dynamics after Ect2-DH addition and found that the temporal shift between peak GTPase activity and F-actin polymerization decreased for both Rho and Cdc42, suggesting that hyperactive Ect2 drives faster signaling between the GTPases and F-actin (Figure 4E-F). We did not observe meaningful changes to any other temporal wave dynamics (Figure S5C-F). Together these results indicate that Ect2 GEF activity increases cortical activation of Rho family GTPases, specifically Rho and Cdc42, and supports faster F-actin assembly. Moreover, our data supports a possible model where Ect2 drives positive feedback signaling for Cdc42, in addition to Rho.

### Rho activity is necessary for Rac and Cdc42 activity and sufficient for excitable F-actin waves

Thus far, our manipulations of known positive and negative feedback regulators of the cortical excitability circuit alter Rho activity, in agreement with previous studies (Bement et al., 2015; Michaud et al., 2022), as well as Cdc42 and Rac. We therefore asked whether Rho activity itself regulates Rac or Cdc42. Previous work has shown that inhibition of Rho activity with C3 exotransferase results in a total loss of cortical F-actin and excitable patterning (Bement et al., 2015; Landino et al., 2021), suggesting that neither Cdc42 nor Rac can compensate for the loss of Rho activity to support cortical F-actin dynamics. Additionally, our observation that Cdc42 and Rac activity peaks after Rho during cortex assembly suggests that Rac and Cdc42 activity is downstream of Rho activation. We therefore hypothesized that Rho activity is the core driver of GTPase activation and F-actin assembly in the artificial cortex.

We tested this idea using C3 exotransferase to inhibit Rho. We again allowed cortex assembly to initiate and performed live addition of vehicle control or 33 μg/mL C3 and assayed GTPase activity over time (Figure. 4A). We observed an immediate loss of Rho in response to C3 treatment, in agreement with previous studies (Bement et al., 2015; Landino et al., 2021; Figure S6A). We also observed a more gradual loss of Rac, Cdc42, and F-actin signal after C3 addition (Figure 5A-B, Figure S6B). This result indicates that Cdc42 and Rac cortical activity is dependent on Rho, which agrees with the observed temporal relationship during cortex assembly. Therefore, we concluded that Rho is the primary driver of cortical assembly and dynamics in artificial cortex.

**Figure 5.**
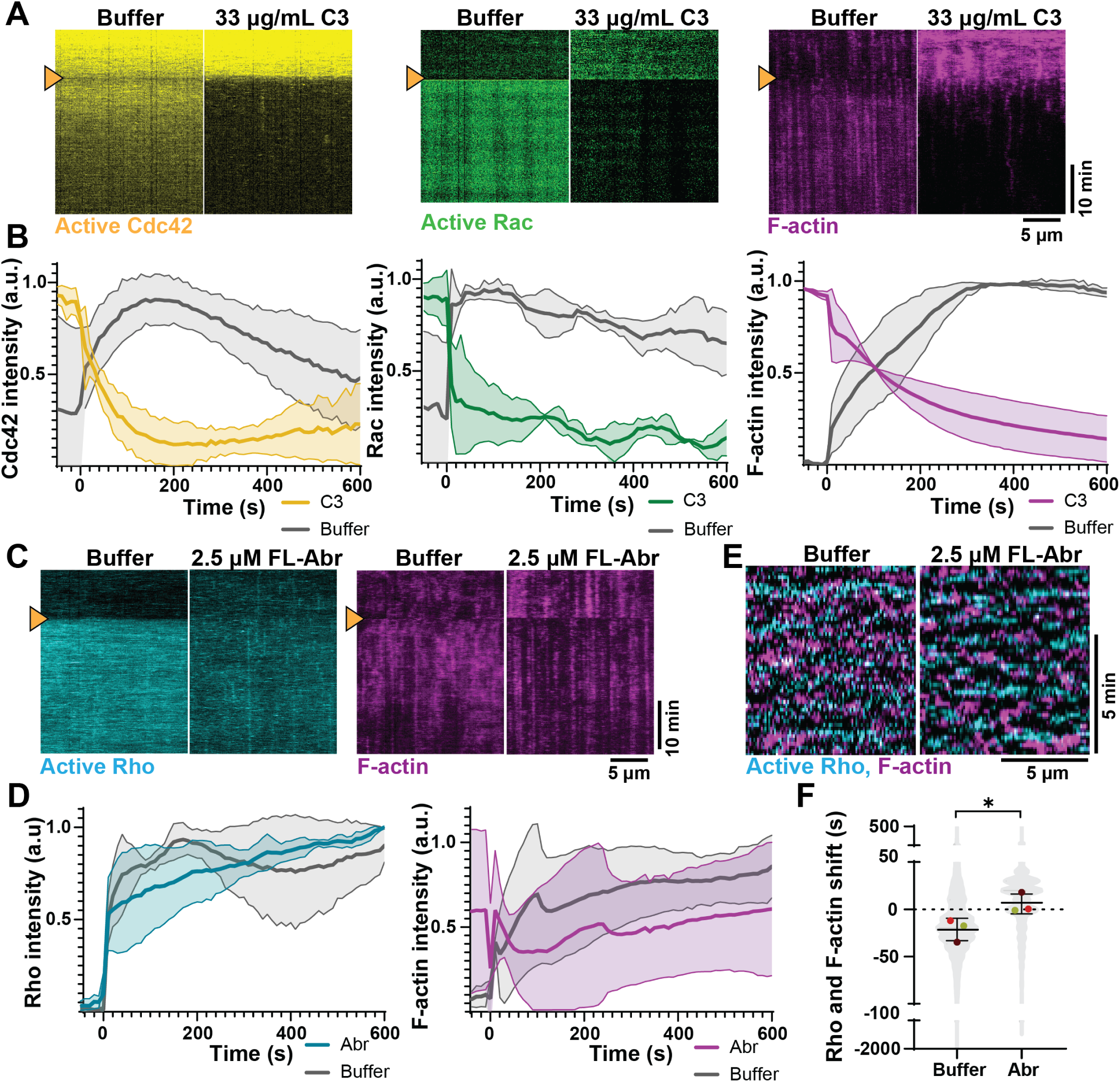
Rho is necessary and sufficient for cortical Rac, Cdc42, and F-actin, while Cdc42 and Rac regulate Rho and F-actin temporal dynamics. **(A)** Representative kymographs of Cdc42, Rac, and F-actin with 33 µg/mL C3 or buffer (vehicle) addition. Intensity is scale-matched across experimental conditions, except for Rac, which exhibits high variability in level of activity. Arrow indicates time of addition. **(B)** Quantification of mean intensity over time of Cdc42, Rac, or F-actin after C3 or buffer treatment. T = 0 represents time of addition. Solid line represents mean, shaded area represents SE. n = 6 movies, 3 extract preps from 3 frogs, 6 ROIs. **(C)** Representative kymographs of Rho and F-actin with Abr or vehicle treatment. Arrow indicates time of addition. **(D)** Quantification of mean intensity over time of Rho or F-actin following FL-Abr or buffer treatment. T = 0 represents time of addition. Solid line represents mean, shaded area represents SE. n = 6 movies, 3 extract preps from 3 frogs, 6 ROIs. **(E)** Kymographs of cortical oscillations of Rho and F-actin after FL-Abr treatment. **(F)** Temporal shift of Rho and F-actin after buffer or FL-Abr treatment. Mean ± SE is shown; violin represents all ROIs analyzed in all movies and dots indicate means of individual experiments. Dot color indicates replicates. n = 6 movies, 3 extract preps from 3 frogs, 4700 ROIs.

### Cdc42 and Rac modulate the temporal relationship between Rho and F-actin

We next asked whether Cdc42 and Rac feedback to Rho activity. To inhibit Cdc42 and Rac in the artificial cortex, we used the dual Cdc42/Rac GAP, Abr (Chuang et al., 1995; Vaughan et al., 2011). We again allowed cortex assembly to initiate and performed live addition of recombinant full-length 2.5 μM Abr or buffer control (Figure S6C, FL-Abr). We found that Abr addition rapidly inhibited Cdc42 and Rac activity in the artificial cortex (Figure S6D-E); however, we did not observe any changes to total Rho or F-actin signal in response to Abr addition (Figure 5C-D). This result suggests that Cdc42 and Rac do not regulate total Rho activation or cortical F-actin assembly.

We further investigated whether Cdc42 and Rac regulate dynamic patterning of Rho and F-actin by quantifying the cortical Rho and F-actin oscillation dynamics after Cdc42/Rac inhibition. We found that the temporal shift between Rho and F-actin decreased after FL-Abr addition (Figure 5E-F). This demonstrates that while Cdc42 and Rac do not regulate total Rho activity, Cdc42 and Rac do feedback into the Rho and F-actin activator-inhibitor circuit, specifically in the maintenance of temporal relationship between Rho and F-actin. This suggests that Cdc42 and Rac have a mechanistic role in regulating Rho and F-actin waves. Taken together, these results demonstrate that Rho activity is necessary and sufficient for cortical F-actin dynamics and that Cdc42 and Rac feedback to Rho and F-actin to drive specific cortical oscillation dynamics.

### Microtubule plus-end overlaps partition cortical Rho and Cdc42 activity

Our findings suggest that Rho family GTPase activity is interdependent and temporally coordinated during cortical patterning. Give the importance of Rho family GTPase spatial organization in other contexts (e.g., wound healing and cell migration) we next investigated spatial organization of GTPase activity in the artificial cortex. Previous work has shown that cortical F-actin waves are spatially regulated at the cytokinetic division plane in *Xenopus* embryos, with decreased oscillation amplitude within the division plane relative to the non-division plane cortex (Swider et al., 2022). One possible mechanism to explain this spatial difference in oscillatory dynamics, is signaling from the microtubule plus-end overlap that develops during anaphase. *In vivo,* microtubule plus-end overlaps support Ect2 accumulation and local Rho activation through directed trafficking of the Centralspindlin complex (Miller and Bement, 2009; Green et al., 2012; Basant and Glotzer, 2018). In interphase egg extract, microtubule plus-end overlaps, organized by artificial centrosomes, also drive cytokinetic signaling, with Rho activity enriched at plus-end overlaps that extend toward the SLB. We therefore investigated GTPase spatial organization in reconstituted 2-dimensional cytokinetic “division planes” (Nguyen et al., 2014; Figure 6A).

**Figure 6.**
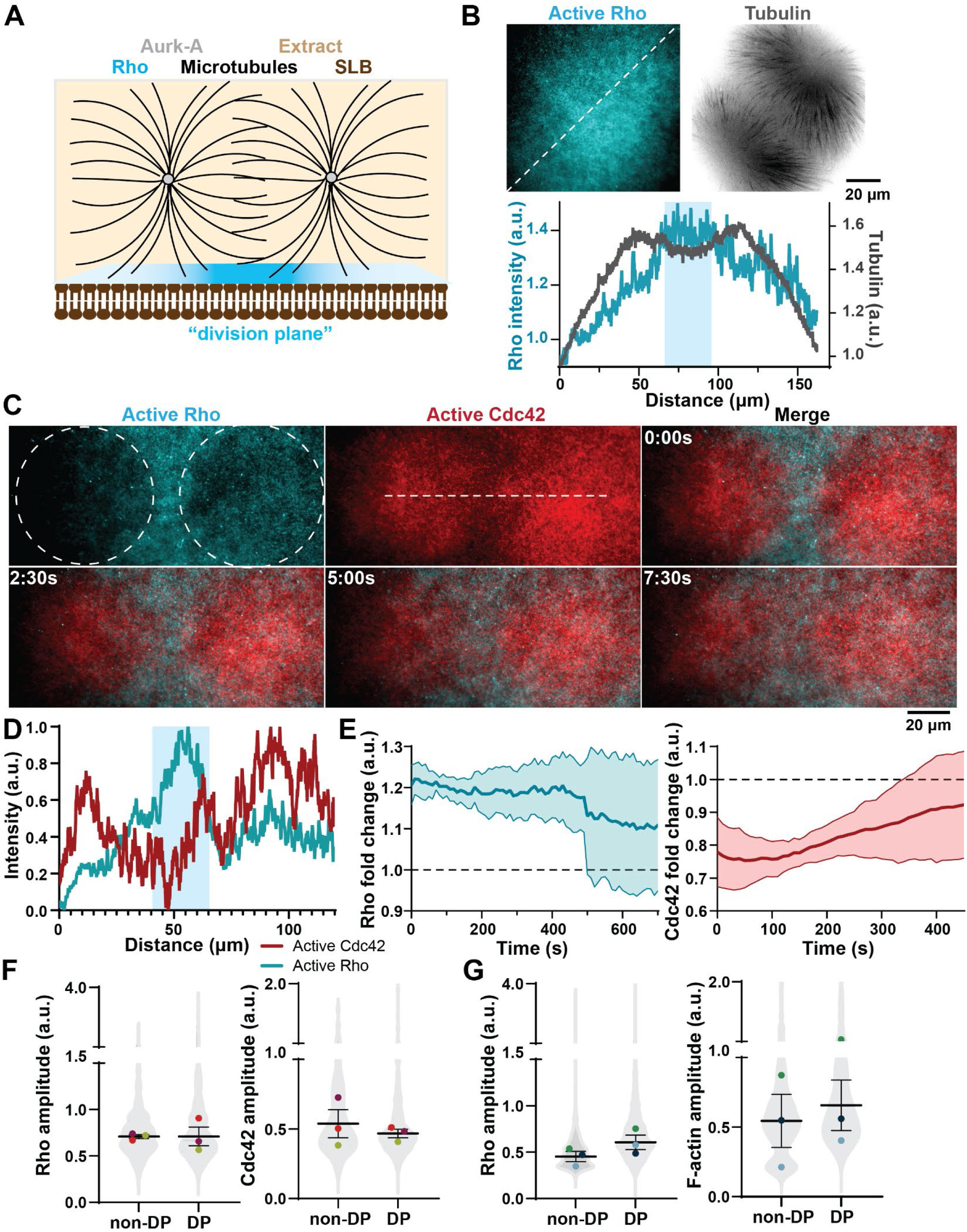
Rho and Cdc42 are partitioned into activity zones by microtubule plus-end overlaps. **(A)** Schematic of AurkA induced microtubule aster formation that drives active Rho enrichment at artificial “division plane”. **(B)** *Top:* Active Rho (cyan) and tubulin (black) organized around anti-Aurora-kinase A beads (artificial centrosomes). Active Rho is enriched at the plus-end overlaps, or “division plane”, between two microtubule asters. *Bottom:* Intensity profile of active Rho and F-actin intensity from representative line shown in A. Blue box indicates distance of the artificial division plane. **(C)** Representative micrographs of Rho and Cdc42 partitioning by microtubule asters. Dotted circles indicate location of microtubule asters. Dotted line indicates region used for intensity profile shown in **(D)**. Time is shown in min:sec. **(D)** Intensity profile of Rho and Cdc42 in the presence of microtubule overlap zones. Blue box indicates “division plane”. **(E)** Quantification of fold change of Rho (left) or Cdc42 (right) at the “division plane” over time. Fold change > 1 indicates increased activity at “division plane”; fold change < 1 indicates decreased activity at “division plane”. Solid line represents mean, shaded area represents SE. n = 3 movies, 3 extract preps from 3 frogs, and 6 ROIs. **(F)** Quantification of Rho and Cdc42 relative amplitude at the division plane (DP) or non-division plane (non-DP) regions. Mean ± SE is shown; violin represents all ROIs analyzed in all movies; dots indicate means of individual experiments. Dot color indicates replicates. n = 3 movies, 3 frogs, 3 extract preps from 1000 ROIs. **(G)** Quantification of Rho and F-actin relative amplitude at the DP or non-DP regions. Mean ± SE is shown; violin represents all regions of interest analyzed and dots indicate means of individual experiments. Dot color indicates replicates. n = 3 movies, 3 frogs, 3 extract preps from 800 ROIs.

In agreement with previous studies, we observed increased Rho and F-actin in the “division plane” (Nguyen et al., 2014; Figure 6A-B, Figure S7A-D). This enrichment of active Rho and F-actin began when microtubule overlaps were first visible within the evanescent field and persisted for approximately 10 minutes across experiments, although F-actin enrichment at “division planes” was more stable than Rho enrichment (Figure 6E, Figure S7C). We next assayed Cdc42 activity in response to microtubule plus end overlaps. We did not assay Rac, because Rac is only active during first 20 min of a reaction (Figure S1A), while microtubule asters take approximately 15-20 minutes to organize (Nguyen et al., 2014). Surprisingly, we found that active Cdc42 is patterned by microtubule plus-end overlaps, but in contrast to Rho, Cdc42 is excluded from the “division plane” and is instead enriched radially, underneath the center of the aster, away from the overlap zone (Figure 6C-D, Figure S7E). This segregation of active Cdc42 and active Rho was transient, with the active Rho and active Cdc42 zones expanding over time (Figure 6C, E).

Given our observation that Rho and Cdc42 activity is interdependent in the artificial cortex in the absence of microtubule plus-end overlaps, we next asked whether this spatial segregation of Rho family GTPases altered cortical oscillation dynamics. We therefore quantified oscillatory dynamics in the reconstituted “division plane” relative to non-division plane regions. Using our image analysis pipeline on defined regions of interest in the field of view, we observed an increase to active Rho and F-actin amplitude compared to non-division plane regions (Figure 6F-G). Additionally, we observed a decrease to active Cdc42 amplitude within the division plane (Figure 6F). This finding demonstrates that cortical Rho, F-actin, and Cdc42 dynamics are differentially regulated in reconstituted cytokinetic zones, organized by microtubule plus end overlaps. Additionally, this data supports a model wherein spatial segregation of Rho and Cdc42 GTPase activity supports regionally distinct cortical oscillation dynamics.

## Discussion

Here we used the artificial cortex system to characterize the spatiotemporal dynamics of Rho family GTPases during cortical patterning. We directly modulated the GTPase activity and regulators of positive and negative feedback to investigate mechanisms of GTPase cross talk. This work has revealed several unique features about the relationship between Rho, Rac, and Cdc42. First, Rho is required for all excitable patterning (Figure 7A), and two pieces of evidence support this model: Rac and Cdc42 activity peak after Rho during cortex assembly, and Rho inhibition abolishes total Rac and Cdc42 activity, even after excitable cortical patterning has been established. Second, Rho family GTPases maintain distinct temporal relationships during cortical patterning. In traveling waves, as well as standing oscillations, Rac activity clearly precedes Rho, Rho precedes Cdc42, and both Rac and Cdc42 act upstream of F-actin (Figure 7A). Notably, we observed differences in the temporal order of GTPase activation in waves (Figures 1 and 2), compared to cortex assembly (Figure S1A). We hypothesize that this difference may indicate that signaling between GTPases during initial cortex assembly differs from the feedback signaling that exists during steady state dynamics. Nevertheless, our data shows that Rho family GTPases maintain a defined temporal activation pattern during excitable behavior, suggesting that unique mechanisms exist to coordinate their spatiotemporal activation and inactivation.

**Figure 7.**
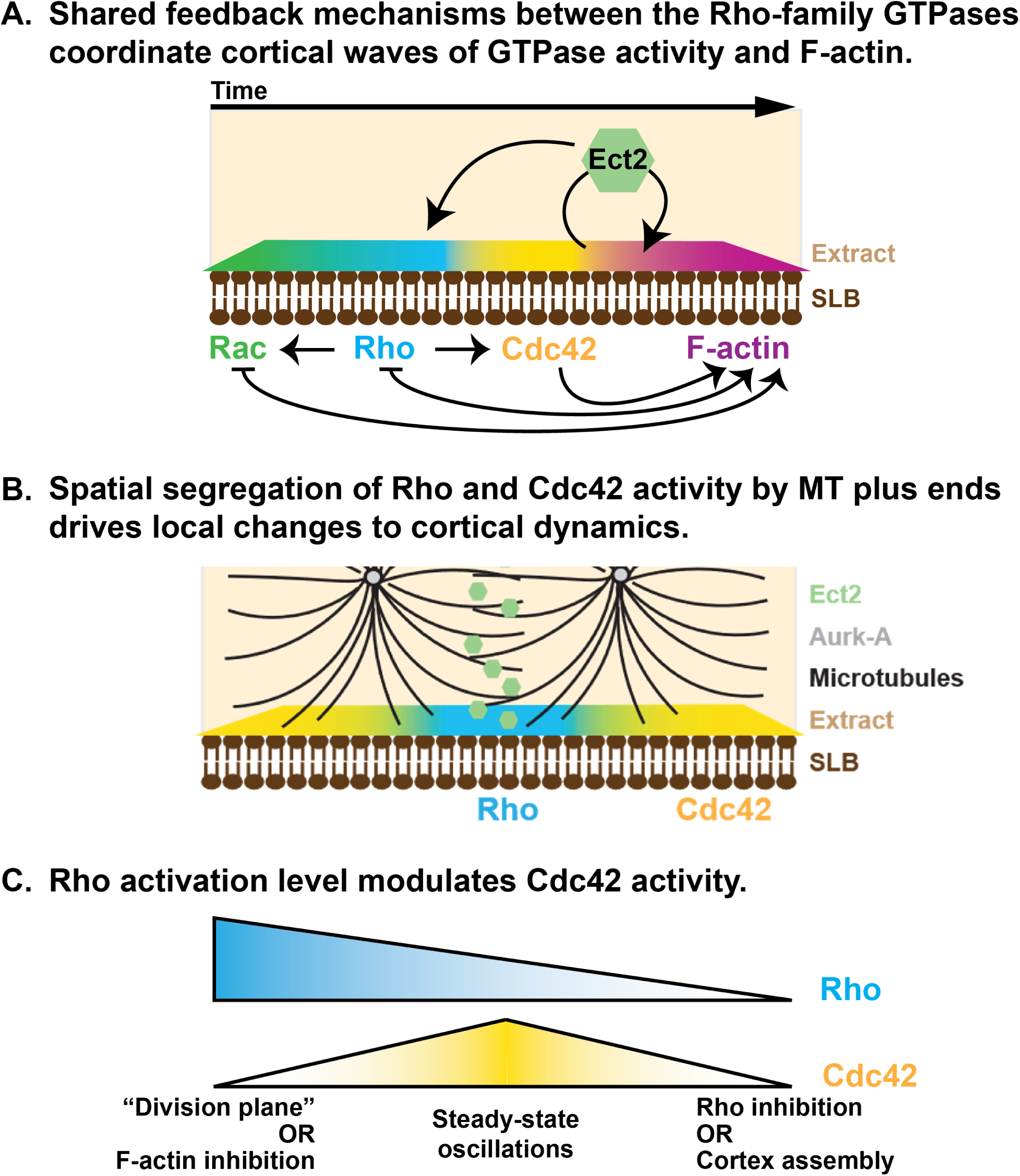
Model of GTPase crosstalk underlying cortical F-actin oscillations and spatial segregation of Rho and Cdc42 GTPases. **(A)** The Rho-family GTPases have a distinct temporal order in excitable wave patterns. Positive feedback from Ect2 amplifies Rho and Cdc42 activity; negative feedback from F-actin inhibits Rho and Rac activity. The Rho-family GTPases regulate each other; Rho activity is necessary for both Cdc42 and Rac activity. **(B)** Microtubule plus-end overlaps are sufficient to induce Rho and Cdc42 activity zones. Cortical F-actin waves dynamics are altered at microtubule plus-ends overlaps. **(C)** Both high and low levels of Rho activity are anti-correlated with Cdc42 activity.

In this study, we present the first evidence that the Rho family GTPases Cdc42 and Rac display excitable behavior in *Xenopus* egg extract. Oscillatory cortical patterning is not unique to Rho; in other systems, Cdc42 waves drive spindle scaling in mast cell division (Fung et al., 2026) and Rac waves coordinate cell migration in T cells (De Belly et al., 2026). Our finding that Cdc42 and Rac participate in traveling waves and standing oscillations indicates they are regulated by coupled positive and negative feedback. As the temporal dynamics of Cdc42 and Rac are matched with Rho and F-actin, mechanisms of feedback may be shared between the GTPases. We find that F-actin dependent negative feedback acts on Rac, and Ect2-dependent positive feedback activates Cdc42 (Figure 7A). As evidence of GTPase crosstalk, we find that Rho is both necessary and sufficient to maintain excitable dynamics of Rac and Cdc42 in the artificial cortex, while Cdc42 and Rac activity feeds back to temporal wave dynamics of Rho and F-actin (Figure 7A). Finally, we find that microtubules partition Cdc42 and Rho into distinct zones, which correlates with spatially distinct F-actin wave dynamics. We propose a model wherein GTPase crosstalk underlies dynamic F-actin patterning, and spatial organization of the Rho family GTPases to drive local changes to cortical oscillation dynamics as have been described for the cytokinetic furrow *in vivo* (Figure 7B).

These studies raise new questions about the mechanisms of positive and negative feedback in regulating excitable dynamics of Rac and Cdc42. We found that while F-actin inhibits Rho and Rac, it may positively regulate Cdc42 as evidenced by the increase in Rac activity and decrease in Cdc42 activity following F-actin inhibition (Figure 7A). Whether F-actin mediates this negative feedback by recruiting a GAP for Rac, as has been shown for Rho (Michaud et al., 2022), is not known. Additionally, Cdc42 and Rac display contrasting responses to F-actin inhibition, suggesting F-actin may recruit regulators that modulate Cdc42 and Rac differently. We also found that Ect2 may provide positive feedback to Cdc42, but not Rac, as we observed that both Rho and Cdc42 activity increases in response to Ect2-DH addition. Importantly, Ect2’s N-terminal domain confers specificity and enables its interactions with other binding partners (Chen et al., 2020). Therefore, additional investigations using full-length Ect2 will be necessary to fully define this relationship.

Furthermore, it remains possible that Cdc42/Rac are only regulated downstream of Rho, and that the observed effects are transmitted through the concomitant changes to Rho activity. For example, increased Cdc42 activity in response to Ect2-DH addition could simply be due to increased Rho. In wound healing and cell migration, Rho respectively activates or inhibits its family members in a spatially specific manner, making this a viable mechanism of Rac and Cdc42 regulation in excitable wave patterning (Vaughan et al., 2011; De Belly et al., 2026). Therefore, Rho activity may be both necessary for and inhibitory to Cdc42 activity (Figure 7C). From our data, we infer that high Rho activity is anti-correlated with Cdc42 activity. This is supported by the observation that Cdc42 activity decreases when F-actin is inhibited and at reconstituted “division planes”. This is a paradoxical result when taken together with our finding that Rho activity is required for Cdc42 activity, as indicated by our observations during cortex assembly and in in response to Rho inhibition by C3 (Figure 7C). This result parallels previous observations in *Xenopus* oocyte wound healing. Basal levels of Rho activity are required to maintain Cdc42 activity, but high levels of Rho activity inhibit Cdc42, enabling segregation of Rho and Cdc42 into activity zones necessary for wound closure (Vaughan et al., 2011).

Crosstalk between the Rho family GTPases has important implications for understanding Rho-mediated actin dynamics during cytokinesis. We investigated this using artificial centrosomes to organize microtubule overlaps zones and 2-D “division planes”. We observed Rho and Cdc42 partition in the presence of plus end overlaps, with Rho enriched at the “division plane” and Cdc42 enriched at the “non-division plane” cortex. This spatial patterning of Rho and Cdc42 may underlie spatially distinct F-actin wave dynamics that we observed in the reconstituted “division plane”. However, our findings in the artificial cortex are distinct from what has been observed *in vivo.* In embryos, F-actin wave amplitude is dampened at the cytokinetic furrow (Swider et al., 2022), whereas we observed a slight increase in F-actin amplitude at the reconstituted “division plane”. This difference may be explained by physiological differences between the early embryo and the artificial cortex system. While the biochemical state of interphase egg extract is analogous to mitotic exit, the artificial cortex system lacks key features of a dividing cell such as chromatin or curved membranes. These features may add additional layers of regulation *in vivo.* Future studies investigating potential additional regulators of GTPase signaling during cytokinesis, like DNA or membrane curvature, will be important for understanding this discrepancy.

## Acknowledgements

We thank the members of the Landino lab for their technical and conceptual feedback on this project. We thank Camilla Bobrowicz for the generation of the Ect2-DH construct used to generate recombinant Ect2-DH protein. We thank Dr. William Bement (University of Wisconsin-Madison) and members of the Bement lab for sharing recombinant protein (full-length Abr) and plasmids (GTPase biosensors, F-actin probe) used in this study and for informing our thinking about cortical dynamics. We thank Dr. Yixian Zheng (Carnegie Institution for Science) for sharing Aurora A antibodies used for generating artificial centrosomes. The Dartmouth Cancer Center Genomics and Molecular Biology Shared Resource provided plasmid sequencing support. Recombinant protein expression was supported by Dartmouth College bioMT core facility through NIH NIGMS grant P20-GM113132.

## Funding

This work was supported by grants from NIGMS awarded to JL (R00GM147826, R35GM166148)

## Author Contributions

Writing – original draft and visualization were performed by AT

Funding acquisition, supervision, and project administration were performed by JL.

Conceptualization, methodology, investigation, and writing – review & editing were performed by AT and JL together.

## Materials and Methods

### Experimental model

Adult, wild-type *Xenopus laevis* females were injected with human chorionic gonadotropin (HCG) to induce egg laying. Frogs received a priming injection of HCG at least 2 days before inducing. Eggs were collected overnight in calcium-free MMR (Ca^2+^-free MMR: 2M NaCl, 40 mM KCl, 20 mM MgCl_2_, 2mM EGTA, 100mM Na-HEPES, pH 7.8).

Frogs were fed twice per week with frog brittle (Nasco or Mazuri LS Aquatic Carni-Blend) and housed in a recirculating tank system (Aquaneering). The water quality parameters (temperature, pH, and conductivity) were constantly monitored for an optimal environment for frog health. Animal care staff monitored the frog’s health and water quality daily.

All studies strictly adhered to the compliance standards of the US Department of Health and Human Services Guide for the Care and Use of Laboratory Animals and were approved by Dartmouth College’s *Animal Care and Use Committee*. A board-certified Laboratory Animal Veterinarian oversees our animal facility.

### Experimental methods

#### Supported lipid bilayers

Powdered lipid stocks (Avanti Polar Lipids: Brain PC, 840053P; Brain PS, 840032P; Liver PI, 840042P; 810150P; Brain PI(4,5)P_2_, 840046P) were resuspended in chloroform (Sigma-Aldrich, 650498) to a 20 mM stock concentration. Lipids were combined in the molar ratios of either 60% PC, 30% PS, and 10% PI (Stock A) or 60% PC, 30% PS, 5% PI, and 5% PI(4,5)P_2_ (Stock B). Lipids were dried down at 42°C with a steady stream of N_2_ gas on a heat block, and then vacuum desiccated for 1 h at 45°C using a speed-vac (Thermo Scientific, SPD1030). Lipid stocks were resuspended in high-salt *Xenopus* buffer (HS-XB: 4 M KCl, 20 mM MgCl_2_, 2 mM CaCl_2_) to 5 mM under N_2_ atmosphere, vortexed, incubated at 37°C for 30 min, and vortexed again. The lipid stocks were sonicated using a cup horn sonicator (QSonica) in polystyrene tubes at 80W for 5 mins with pulsed sonication (30s on; 10s off) to create small unilamellar vesicles (SUVs) under N_2_ atmosphere at room temperature. SUVs were filtered using a 0.22 µm filter (Fisher Scientific, SCGP00525) into borosilicate glass vials (Thermo Scientific, 4010-17) under N_2_ atmosphere and stored in −20°C for less than 3 months.

SUVs were thawed at room temperature under N_2_ atmosphere to create supported lipid bilayers (SLBs). Stock A and Stock B lipid mixes were combined at the appropriate ratio to generate 0.1% PIP_2_ SLBs. SUV solutions were sonicated under N_2_ atmosphere and diluted to 0.5 mM using HS-XB, and CaCl_2_ was added to 4 mM. CaCl_2_-treated SUVs were added to O_2_ plasma-cleaned imaging wells and incubated at 37°C for 30 min. The SLBs were washed three times with *Xenopus* buffer (XB: 100 mM KCl, 1mM MgCl_2_, 10 mM HEPES, pH 7.7) without exposing the SLBs to air. Washed SLBs were kept at room temperature for less than 4 h before imaging experiments.

#### Preparing imaging wells

Coverslips (24 x 50 mm No. 1.5, VWR) were washed in a solution of 2% Hellmanex III (Sigma) for 2 hours at 70°C with constant stirring. Coverslips were rinsed thoroughly with deionized water and allowed to dry overnight in a dust-free chamber until assembly of imaging chambers. A cleaned coverslip was attached to a custom aluminum coverslip mount, comprised of a rectangular slide with a central oval-shaped opening and clamps on either end to hold the coverslip in place (Landino et al., 2021; University of Michigan, Scientific Instrument Shop). The cap and lower third of 0.2 mL PCR tubes (Sigma) were removed to create wells for imaging extract. The rim of the PCR tube was dotted with UV glue (Norland Optical Adhesive No. 61, ThermoFisher Scientific) and adhered to the coverslip using 365 nM UV light for one minute. Additional glue was used to reinforce the attachment of the wells, and chambers were exposed to 365 nM UV light for 30 minutes. Chambers were stored in a dust-free container at room temperature until use.

Immediately prior to adding SUVs, imaging chambers were O_2_ plasma cleaned (PE25-JW Benchtop Plasma Cleaning System, PLASMA ETCH) for 10 minutes.

#### Actin intact extract prep (CSF-arrested, M-phase)

Adult, female *X. laevis* were induced with HCG and housed in Ca^2+^ free Marc’s Modified Ringers (Ca^2+^-free MMR: 100 mM NaCl, 2mM KCl, 1mM MgCl_2_, 100 µM EGTA, 5mM HEPES, pH 7.8) to lay eggs. Eggs were de-jellied with 2% cysteine (Sigma-Aldrich, 168149) in Ca^2+^-free MMR at pH 7.8 for 10 min or until the egg’s jelly coat was removed. The eggs were washed twice with cytostatic factor *Xenopus* buffer (CSF-XB: XB + 1 µM MgCl_2_, 5 mM K-EGTA) while removing lysed and activated eggs (gardening). De-jellied eggs were washed in CSF-XB with a protease inhibitor cocktail (LPC: 10 µg/mL Leupeptin, Pepstatin A, and Chymostatin; Sigma-Aldrich, L2884, 11524488001, C7268) and transferred to a thin-walled UltraClear centrifuge tube (Beckman Coulter) submerged in CSF-XB + LPC. The eggs were packed at 700 x g for 30 s followed by 1,400 x g for 15 s in a clinical centrifuge (Internation Equipment Co. Model CL). Excess buffer was removed after packing. The eggs were crushed in an ultracentrifuge (Beckman) using a swinging bucket rotor (SW55) at 15,000 x g for 15 min at 4°C. The cytoplasmic fraction was harvested using a needle and syringe (Fisher, 14955464; BD, 305196) through the side of the centrifuge tube while not disturbing the lipid or pellet fractions. Protease inhibitors (LPC) were added to a final concentration of 10 µg/mL with an energy mix with a final concentration of 9 mM (energy mix: creatine phosphate, 1mM ATP, 1mM MgCl_2_; Sigma, CRPHO, M8266). The extract is arrested in metaphase of meiosis II and kept on ice until use.

#### Converting to I-phase and contractility assay

To confirm high-quality M-phase arrest, 2 µl droplets of the extract were added to 2 ml of mineral oil (Crystalgen, 24-107) in 35mm well dishes (Fisher Scientific, 3535) and monitored under a stereoscope (VWR, 89404-492). High-quality extract was visibly contracted in the mineral oil after 5 min. Noncontractile egg extract was not used. CaCl_2_ (Fisher Scientific, AC349610000) was added to extract to a final concentration of 0.4 mM, vigorously mixed, and incubated at room temperature for 5 min. Extract was put back on ice to prevent further progression through the cell cycle and was assayed for contractility. The successfully cycled I-phase extract was not contractile after 5 min under oil.

#### Freezing extract

Egg extract was gradually frozen in microfuge tubes in a cryofreezing chamber (Analytics Shop, NL51000001) in a −80°C freezer overnight.

#### Preparing reactions for imaging

Recombinant protein probes were added to egg extract at the final concentrations: GFP-rGBD or mCherry-rGBD, 750 nM, GFP-pGBD, 300-750 nM, mCherry-wGBD, 750nM, UtrCH-594 or UtrCH-647, 200 nM, AF647-Tubulin (Cytoskeleton, TL670M) 400-1000 nM. Extract with recombinant probes was added to washed SLBs immediately transferred to the microscope for imaging.

### Live imaging

For all experiments described here, total internal reflection fluorescence (TIRF) microscopy was performed using a Nikon iLas Ring TIRF, with a Ti2 inverted, motorized microscope base, LUNF laser launch (488 nm/561 nm/640 nm), perfect focus, and photo-stimulation module controlled by NIS Elements software. A TIRF Quad Dichroic cube (C-FL TIRF Ultra Hi S/N 405/488/561/638 Quad Cube, Z Quad HC Cleanup, HC TIRF Quad Dichroic, in metal cube, HC Quad barrier Filter) was used with a 60X Objective lens (CFI60 Apochromat TIRF 60X Oil Immersion Objective Lens N.A. 1.49, W.D. 0.12 mm, F.O.V 22 mm) and a Photometrics Prime 95B Back-illuminated sCMOS camera.

For investigating active Rho and F-actin wave dynamics, imaging started within 5 min of the addition of extract with the recombinant protein probes to the SLBs. The SLBs were imaged in 10s intervals for 30–60 min in the appropriate channels with perfect focus for the duration of the live imaging.

For cortex assembly assays, imaging wells were imaged on the TIRF microscope without extract (the first 2-5 frames of videos). Extract with recombinant probes was added to the well while acquiring images and imaged for 10 minutes at 2s intervals.

### Drug treatments and perturbations

*Latrunculin A powder* (Tocris Bioscience, 3973) was resuspended in DMSO to a working concentration of 300 μM and stored at -20°C. Egg extract was treated with 15 μM Lat A, or an equivalent volume of DMSO as a vehicle control for 5 min at room temperature before adding to SLBs.

*Lyophilized C3 transferase* (Cytoskeleton, CT03) was resuspended in buffer containing: 500 mM imidazole, 50 mM TrisHCl, pH 7.5, 10 mM MgCl_2_, 200 mM NaCl, 5% sucrose, 1% dextran and supplemented with 1 mM DTT. C3 transferase was aliquoted, frozen in liquid N_2_, and stored at −80°C until use.

*For real-time addition of C3 transferase, Ect2-DH, or Abr*, an equivalent volume of I-phase extract containing 2X C3 transferase (final concentration of 33 μg/mL), Ect2-DH (final concentration of 100 nM), Abr (final concentration of 2.5 µM) or buffer (1X XB) was added to the imaging chamber at the time indicated. Extract containing 2X recombinant protein or buffer was kept at room temperature for no more than one minute prior to addition to the imaging chamber.

### Anti-AurkA beads and microtubule asters

Anti-AurkA beads were prepared as previously described (Tsai and Zheng, 2005; Field et al., 2017). Beads were added to egg extract with fluorescent probes prior to addition to the bilayer. Microtubule asters took 10-20 minutes to self-organize around beads, which agrees with previous descriptions of microtubule aster assembly in egg extract (Tsai and Zheng, 2005; Field et al., 2017). Fields of view that contained two or more microtubule asters were selected for imaging plus end overlaps.

### Generation of new DNA constructs

DNA constructs for insect cell expression of mCherry-wGBD, mCherry-rGBD, and GFP-pGBD were generated by Gibson assembly into a pFastBac-derived expression vectors. Inserts were amplified by PCR from Addgene plasmids 26734, 26732, and 26735, respectively.

pFastBac1/FLAG-Ect2-DH was generated from *Xenopus* Ect2 (Bement et al., 2015), and amino acids 415-884, which contain the DH (GEF activity) and PH (membrane binding) domains (Kanada et al., 2008), were amplified using PCR. Gibson assembly was used to incorporate them into a pFastBac expression vectors. The amino acid sequence used for generating Ect2-DH is as follows:

SLLDISNTPESGKSFSDTPKIIKPSKVSTPVPQQSARWQVAMELYQTESNYVDILTTIVQLFQLPLEKEGQLGGPILAQEEIKTIFGSIPDILDVHTNIKGDLEKLMIDWAESKSIGDIILTYSKDLEKIYPPFVNFFEMSKETIIKCEKQKPRFHAFLKINQSKPECGRQTLVELLIRPVQRLPSVALLLNDIKKHTADDNPDKVLLEKAIDSLKQVMTYINEDKRKTEGQKQIFDVVYEVDGCPANLLSSHRMLVQRVETIALGEDLCDRGEQVTLFLFNDCLEIARKRHKGIGAFKSPHGHTRPPACLKHICLMPLSQIKKVLNVKDTEECHNAFALVVRPPTEQTNQLFSFQLTTEELPKVDWLKMLCRHVANTICKADAENLIYSTDADSLEVSTKDVDSTLSRASRAIKKTS KKVTRAFSFTKTPKRALQRALMVQNADGRSPGPSNEGFASCRMPSTSSL

All constructs were verified by sequencing.

### Expression and purification of recombinant protein

*Active Rho, Cdc42, and F-actin probes; Ect2-DH and Abr proteins*. For recombinant protein expression of rGBD, wGBD, UtrCH, and Ect2-DH, pFastBac1 constructs encoding FLAG-GFP-rGBD, 6xHis-GFP-pGBD, FLAG-mCherry-wGBD, FLAG-Cys-UtrCH, and FLAG-Ect2DH were used to generate recombinant bacmids in DH10BAC (Invitrogen) bacteria. Sf9 infected cells were transfected with recombinant bacmids with PEI transfection reagent (Polysciences, 3966) in collaboration with the bioMT core facility at Dartmouth. The virus was amplified to improve titer, and Sf9 cells were infected for three days at 27°C for protein expression.

For FLAG purifications, cell pellets were lysed in PBS (pH 7.8) + 0.1% TX-100 with protease inhibitors (LPC, PMSF, Sigma) and solubilized with end-over-end mixing for 1 h at 4°C. The solubilized sample was centrifuged for 10 min, approximately 20K × g, at 4°C. Recombinant proteins were purified from the supernatant fraction using anti-FLAG M2 affinity resin (Sigma), equilibrated with lysis buffer. The protein-bound resin was washed with 25 mM HEPES, pH 7.5, 100 mM KCl, and protease inhibitors. Protein was eluted using arginine-based elution, 1 M Arginine, pH 4.4 (Bement et al., 2015). Elution fractions were collected in an equivalent volume wash buffer.

For His purifications, cell pellets were lysed in 2x PBS (pH 8.0), 20 mM Imidazole, (pH 7.4), 500 mM NaCl, 0.1% Triton X-100, and 14 mM BME with protease inhibitors (LPC, PMSF, Sigma) and solubilized with end-over-end mixing for 1 h at 4°C. The solubilized sample was centrifuged for 10 min, approximately 20K × g, at 4°C. Recombinant proteins were purified from the supernatant fraction using Ni-NTA affinity resin (Sigma), equilibrated with lysis buffer. The protein-bound resin was washed with 1x PBS (pH 8.0), 20 mM Imidazole (pH 7.4), 500 mM NaCl, 0.1% Triton X-100, and 14 mM BME. Protein was eluted using imidazole-based elution: 1x PBS (pH 8.0), 300 mM Imidazole (pH 7.4), 150 mM NaCl, 0.1% Triton X-100, and 14 mM BME.

Purity of recombinant protein in fractions was evaluated by SDS-PAGE. Fractions were pooled and protein was concentrated using Amicon Ultra Centrifugal Filters, 30 kDa MWCO (Millipore) and stored in 25 mM HEPES, 100 mM KCl, pH 7.5. FLAG-Cys-UtrCH was labeled using Alexa Fluor 594 Protein Labeling Kit or Alexa Fluor 647 Protein Labeling Kit targeting primary amines according to the manufacturer’s instructions (ThermoFisher). Protein was frozen using liquid N_2_ and stored at −80°C.

FLAG-Abr was a gift from Bill Bement (University of Wisconsin, Madison, WI) and described in (Vaughan et al., 2011; Hoachlander-Hobby et al., 2026).

### Quantification and statistical analysis

#### Cortex assembly assay

To measure the GTPase activation and F-actin polymerization during cortex assembly, mean intensities were normalized between 0 and 1 by subtracting the minimum intensity from all values, and subsequently dividing by the maximum intensity from all values, using the following equation.

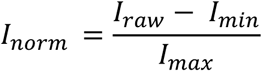

Where *I_raw_* is the mean gray value at any given time point, and *I_min_* is the minimum of all timepoints, and *I_max_* is the maximum of all timepoints calculated after *I_min_* has been subtracted from *I_raw_* at each timepoint.

#### Quantification of whole-field intensity of GTPases and F-actin in response to drug treatment or other perturbations

Whole-field intensity was measured using the raw data (not difference subtracted), cropped to the central portion of the field of view (125 µm × 125 µm). Mean gray values were measured using Fiji “Multi Measure” tool. For LatA treatment (Figure 3, Figure S4), mean gray values were normalized to the mean intensity of all time points for each channel and each condition. Then, the fold change calculation was performed by dividing mean intensity by the average mean gray value from the last 10 frames of the movie, using the following equation.

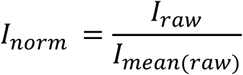

Where *I_mean_*_(*raw*)_ is the average of the mean gray value of all frames for the movie.

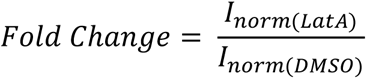

Where *I_norm_*_(*LatA*)_ and *I_norm_*_(*DMSO*)_ are the average of 10 frames at 20 minutes (for Rac movies) or 60 minutes (for Cdc42 movies) for each experimental condition.

For C3 and Abr addition experiments (Figure 5, Figure S6), mean intensities were normalized between 0 and 1 by subtracting the minimum intensity, and subsequently dividing by the maximum intensity, using the equation described for the cortex assembly assay. For Ect2-DH addition (Figure 4, Figure S5) experiments, mean intensities were normalized to the average mean intensity of the frames prior to Ect2-DH addition for each movie, using the following equation.

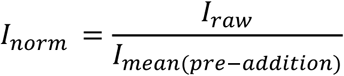

Where *I_mean_*_(*pre*−*addition*)_ is the average of the mean gray values prior to Ect2-DH addition.

For all experiments, normalized mean intensity or fold change was plotted in Prism (GraphPad).

#### Image pre-processing for analyzing wave dynamics

Videos of active Rho and F-actin dynamics on SLBs were processed in Fiji (Schindelin et al., 2012) and cropped to the central portion of the field of view at approximately 150 µm × 150 µm. Relevant frames were selected for analysis based on the visible appearance of the active Rho signal. Each channel was adjusted for visualization and pseudo-colored as indicated in the figures. To remove a non-dynamic signal, difference subtraction was performed by duplicating the time series stacks with a 30 s time difference and using the “image subtraction” tool in Fiji as previously described (Bement et al., 2015; Landino et al., 2021). Kymographs were created using the “Multi Kymograph” tool in Fiji from a line of width 15 pixels drawn across the video, as indicated in the figures. Kymographs were created from each channel individually and then merged.

#### Analysis of active Rho family GTPases and F-actin wave dynamics

Difference-subtracted videos were analyzed using *waveanalysis* (Landino et al., 2021; Swider et al., 2022) to quantify dynamics of active GTPases and F-actin over time. The field of view was subdivided into boxes 4 µm × 4 µm in size, resulting in thousands of boxes (bins) from a single movie. Data were thresholded for a cross-correlation function that indicated active Rho signal leads F-actin (negative peak values, Figure S2D-E; Figure S3A-B). For standing oscillation analysis, data were also thresholded for an auto-correlation function that was centered around 1 (Figure S3C). For Active Rac analysis, movies were qualitatively screened for oscillatory Rac activity (Figure S1B). Movies or timepoints not displaying Rac oscillations were not used for *waveanalysis*. Largely, oscillatory Rac activity was only observed in the first 30 minutes of the extract reaction. Wave dynamics for each bin as well as means for each movie were plotted in Prism (GraphPad).

#### Microtubule aster experiments

Areas of active Rho enrichment (“division planes”) between asters were qualitatively determined by eye and cropped. These areas ranged 15 - 45 μm wide and 70 - 100 µm long, which agrees with previous literature. For each experiment, control ROIs of an equivalent area were analyzed directly underneath asters of “non-division plane” zones. Difference subtraction and *waveanalysis* were subsequently performed on these zones. For movies wherein multiple “division planes” were visible, multiple crops of “division planes” and non-“division planes” were generated and used for *waveanalysis*. *Waveanalysis* values for each zone within one movie were averaged and treated as a single replicate.

For quantification of GTPase and F-actin fold change, an approximately 20 x 20 μm box was drawn within each of these zones, and mean gray values were measured for the respective channels using the “multi-measure” tool in Fiji. Fold change was calculated by dividing the mean gray value within the “division plane” by the mean gray value at the non-“division plane” for each timepoint. Quantification GTPase and F-actin patterning by microtubule asters was performed using the “Plot Profile” tool in Fiji. Lines were drawn by eye (between 15-20 pixels wide), approximately halfway through the width of the microtubule aster, perpendicular to the area of active Rho enrichment.

### Statistical Analysis

Welch’s t-test was used to determine the statistical significance of Rho and F-actin temporal shift after Abr treatment.

## Abbreviations

*X. laevis*: *Xenopus laevis*
F-actin: Filamentous actin
GTPase: Guanine tri-phosphatase
SLB: Supported Lipid Bilayer
GEF: Guanine nucleotide Exchange Factor
GAP: GTPase-Activating Protein
TIRF: Total Internal Reflection Fluorescence
GFP: Green Fluorescent Protein
rGBD: Rhotekin GTPase binding domain
UtrCH: Utrophin calponin homology domain
wGBD: N-WASP GTPase binding domain
pGBD: Rac GTPase binding domain of Rhotekin
LatA: Latrunculin A
DH: *Dbl* homology domain
PH: Pleckstrin homology domain
C3: C3 exoenzyme, *clostridium botulinum*
DP: Division plane
PC: Phosphatidylcholine
PS: Phosphatidylserine
PI: Phosphatidylinositol
PIP_2_: Phosphatidylinositol 4,5-bisphosphate

## Notes

### Competing Interest Statement

The authors have declared no competing interest.

