## Supplemental Files for "Rho family GTPase crosstalk through shared positive and negative feedback mechanisms underlies cytokinetic F-actin waves"

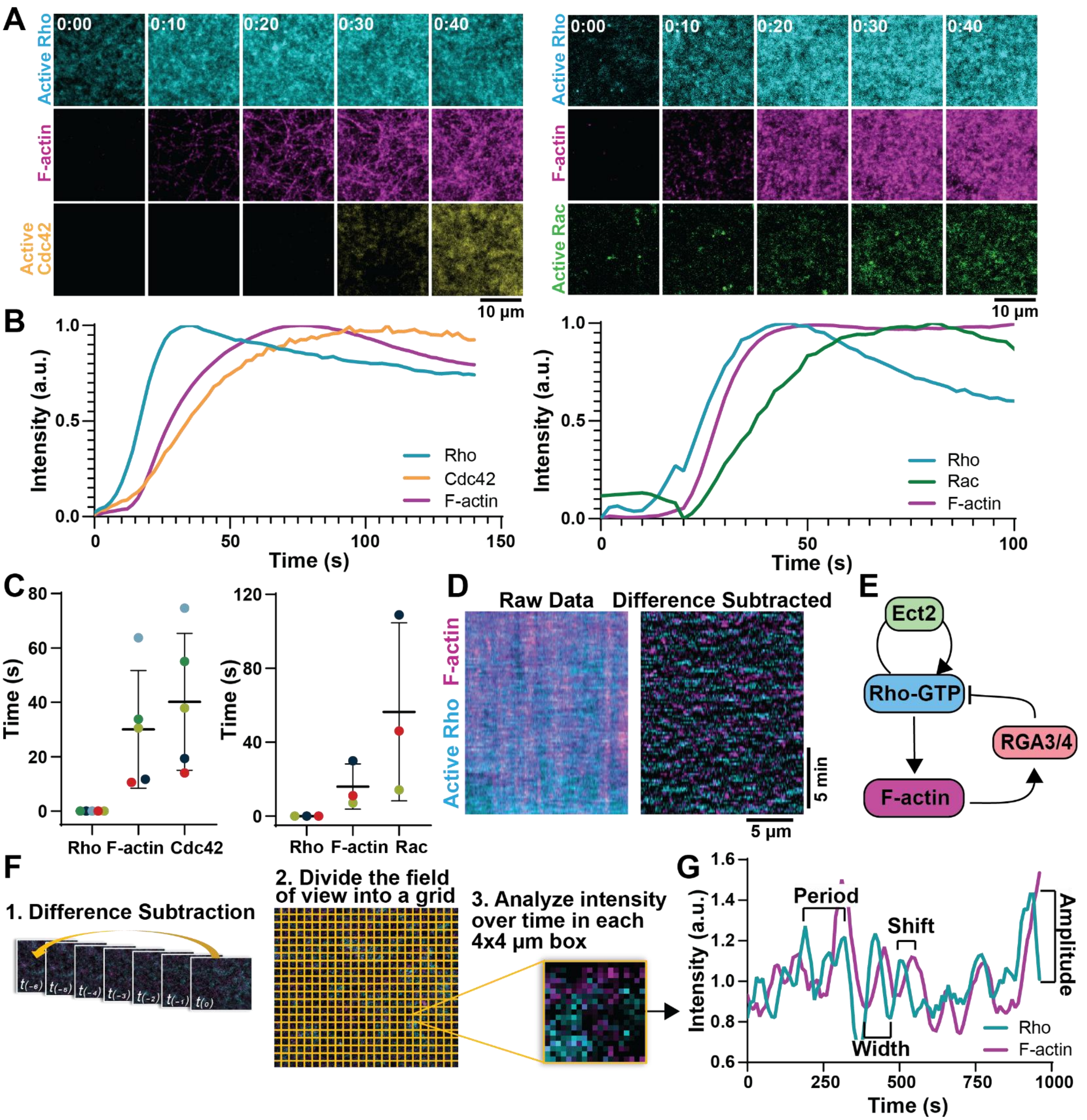


**Supplemental Figure 1. Molecular circuit and analysis pipeline for F-actin cortical waves.**

**(A)** *Left:* Representative micrographs of extract addition to naïve bilayer with probes for active Rho and Cdc42 or *Right:* active Rho and Rac. Time is indicated in min:sec. **(B)** Representative intensity over time plots for extract addition to naïve bilayer assay for Rho, Cdc42, and F-actin (left) or Rho, Rac, and F-actin (right). **(C)** Time to half-max intensity for GTPase activation and F-actin assembly. Mean ± standard error (SE) is shown; dots indicate individual experiments. For Cdc42: n = 5 movies, 5 extract preps from 5 frogs, 5 ROIs. For Rac: n = 3 movies, 3 extract preps from 3 frogs, 3 ROIs. **(D)** Example kymograph of Rho and F-actin oscillations before (raw data, left) and after temporal difference subtraction (right). **(E)** Fast positive feedback (Ect2) and delayed negative feedback (RGA3/4) underlies Rho and F-actin wave dyanmics. **(F)** *Waveanalysis* pipeline (Swider et al., 2022). Temporal difference subtraction is performed on live imaging data to remove non-dynamic signal. Custom code is used to plot intensity over time for 4 x 4 μm bins. **(G)** Intensity traces are used to quantify wave dynamics for each bin including temporal period, temporal width, temporal shift and wave amplitude. 1 bin = 1 ROI.

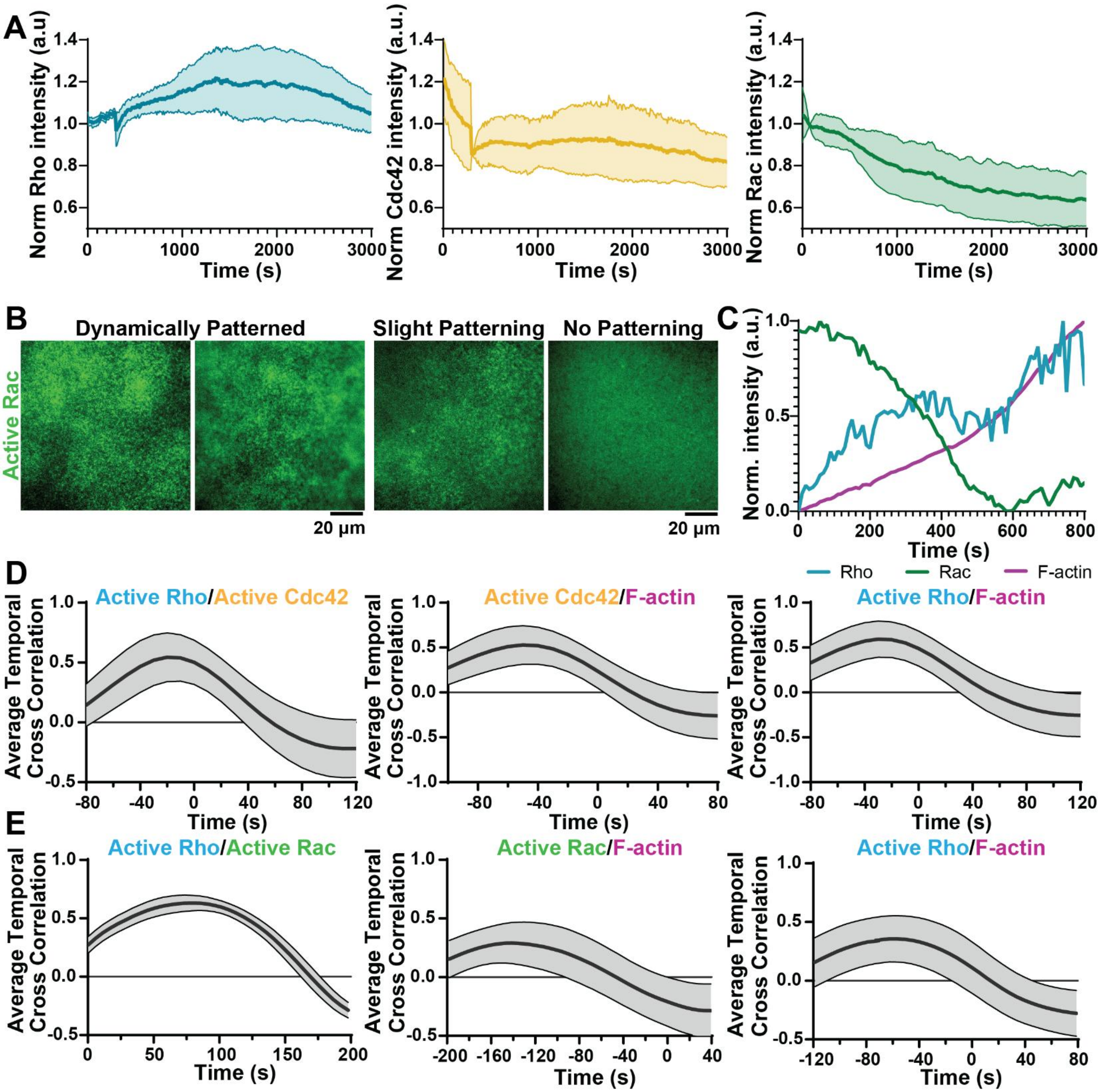


**Supplemental Figure 2. GTPase activity over time and traveling wave cross-correlations.**

**(A)** Total Rho, Rac and Cdc42 intensity over time. Rho activity stays constant; Rac and Cdc42 decrease over the couse of 1 hour. Solid line indicates mean, shaded area indicates SE. n = 3 movies, 3 extract preps from 3 frogs, 3 ROIs. **(B)** Represenative micrographs of Rac dynamic patterning across SLB. Left examples were scored as "dynamically patterned" and used for quantification of wave dynamics. Right examples were not. **(C)** Representative example of anti-correlated Rac vs Rho and F-actin mean intensities over time. **(D)** Representative cross-correlation functions for Rho, Cdc42, and F-actin in traveling waves from one movie. Black line represents mean cross-correlation, gray shading indicates standard deviation (SD). **(E)** Representative cross-correlation functions for Rho, Rac, and F-actin in traveling waves from one movie. Solid line represents mean cross-correlation, shaded area indicates SD.

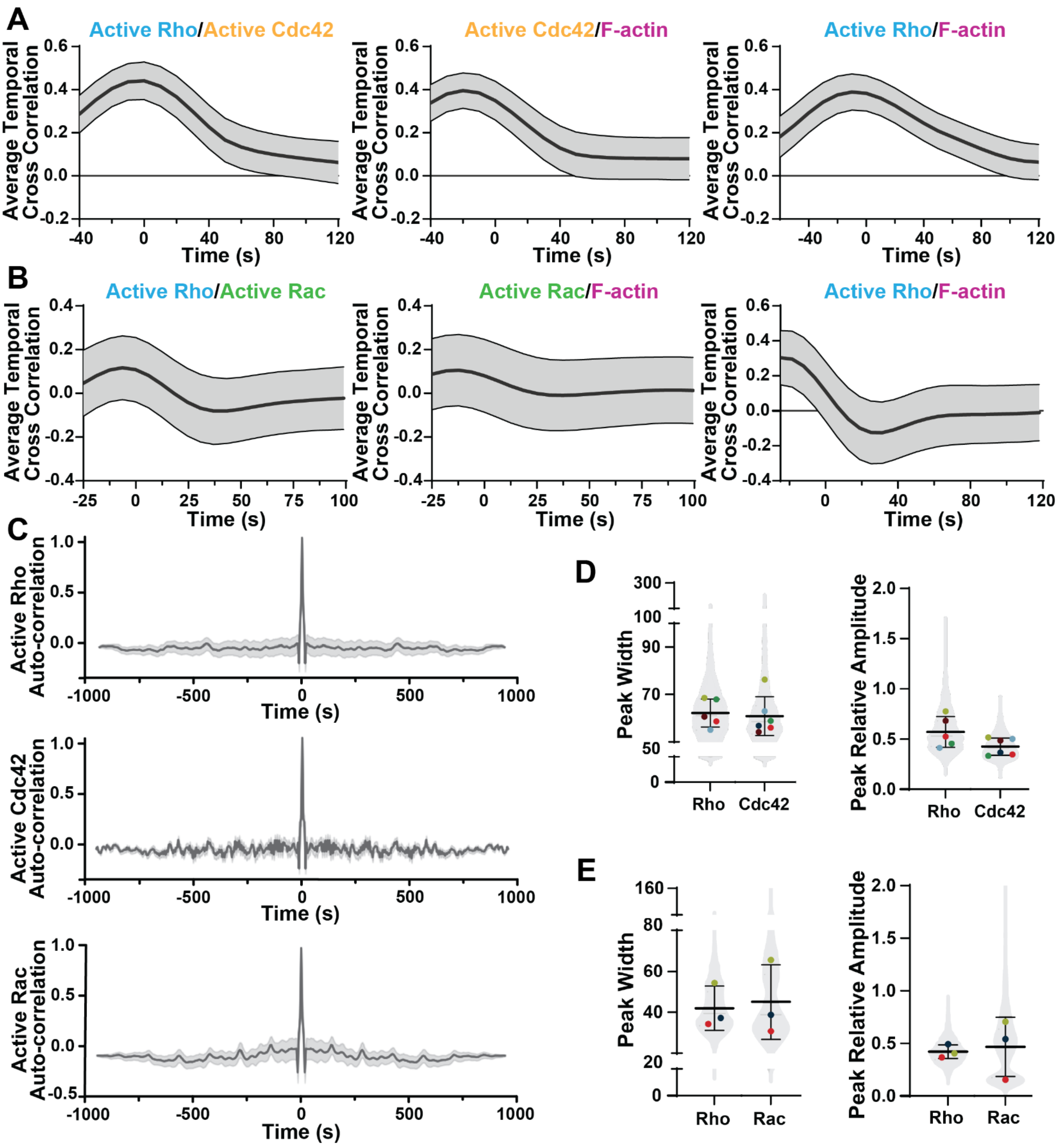


**Supplemental Figure 3. GTPase oscillation cross-correlations and wave dynamics.**

**(A)** Representative cross-correlation functions for Rho, Cdc42, and F-actin in oscillations from one movie. Solid line represents mean cross-correlation, shaded area indicates SD. **(B)** Representative cross-correlation functions for Rho, Cdc42, and F-actin in oscillations. Black line represents mean cross-correlation, gray shading indicates SD. **(C)** Representative auto-correlation functions for Rho, Cdc42, and Rac. Black line represents mean cross-correlation, gray shading indicates SD. **(D)** Quantification of relative amplitude and peak width of Rho, Cdc42, and F-actin. Mean ± SE is shown;

violin represents all ROIs analyzed for all movies; dots indicate means of individual experiments. Dot color indicates replicates. n = 6 movies, 6 extract preps from 6 frogs, 4698 ROIs. **(E)** Quantification of relative amplitude and peak width of Rho, Rac, and F-actin. Mean ± SE is shown; violin represents all ROIs analyzed for all movies; dots indicate means of individual experiments. Dot color indicates replicates. n = 3 movies, 3 extract preps from 3 frogs, 2350 ROIs.

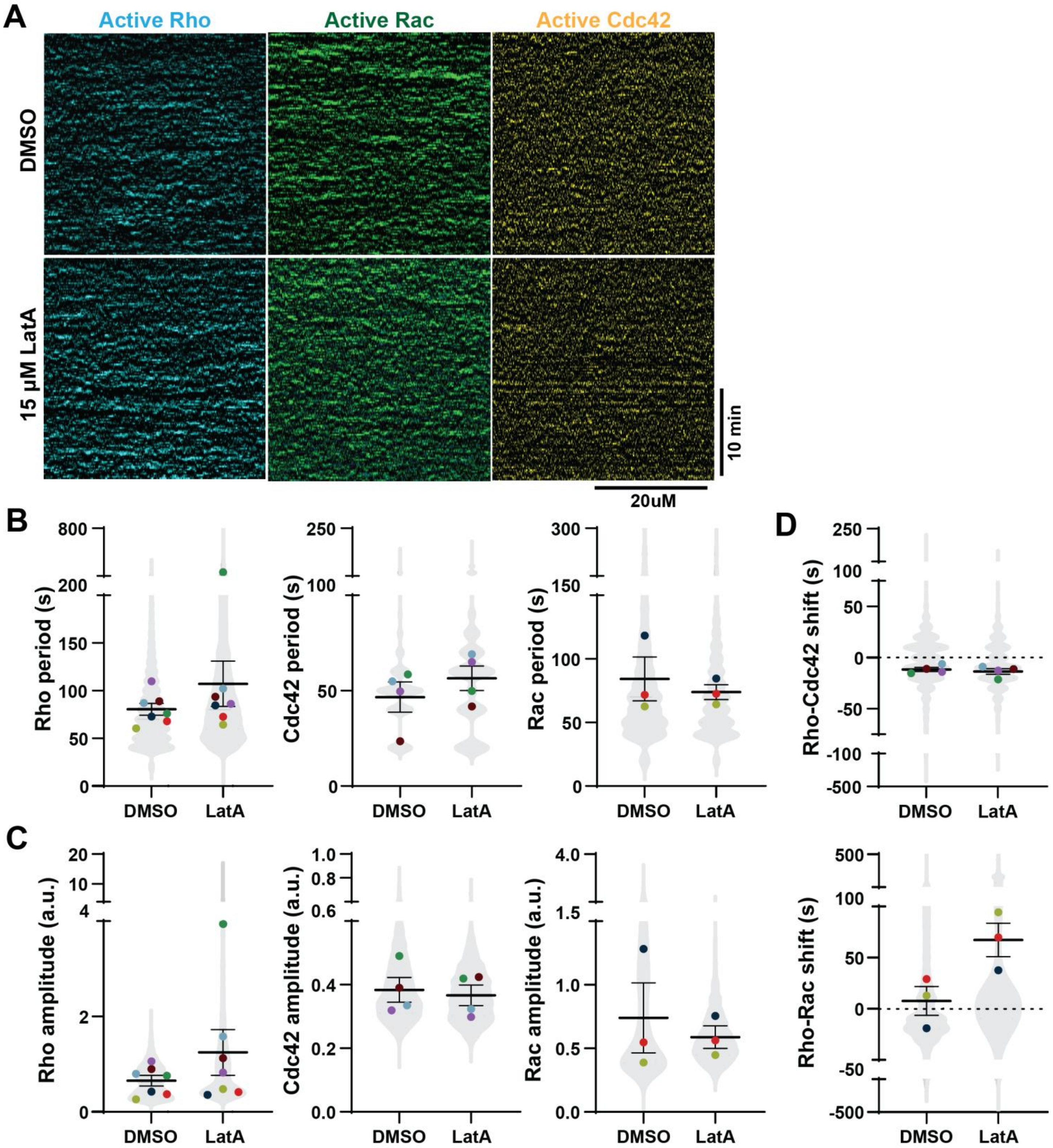


**Supplemental Figure 4. Wave dynamics following LatA treatment. (A)** Representative kymographs of GTPase oscillations after DMSO or 15 µM LatA treatment. **(B-C)** Quantification of GTPase period and amplitude following DMSO or 15 µM LatA treatment. Mean ± SE is shown; violin represents all ROIs analyzed for all movies; dots indicate means of individual experiments. Dot color indicates replicates. Rho: n = 14 movies, 7 extract preps from 7 frogs, 10,322 ROIs**.** Cdc42: n = 8 movies, 4 extract preps from 4 frogs, 5632 ROIs. Rac: n = 6 movies, 3 extract preps from 3 frogs, 7400 ROIs. **(D)** Quantification of temporal shift after DMSO or 15 µM LatA treatment. Mean ± SE is shown; violin represents all ROIs analyzed for all movies; dots indicate means of individual experiments. Dot color indicates replicates. Cdc42: n = 8 movies, 4 extract preps from 4 frogs, 5632 ROIs. Rac: n = 6 movies, 3 extract preps from 3 frogs, 4700 ROIs.

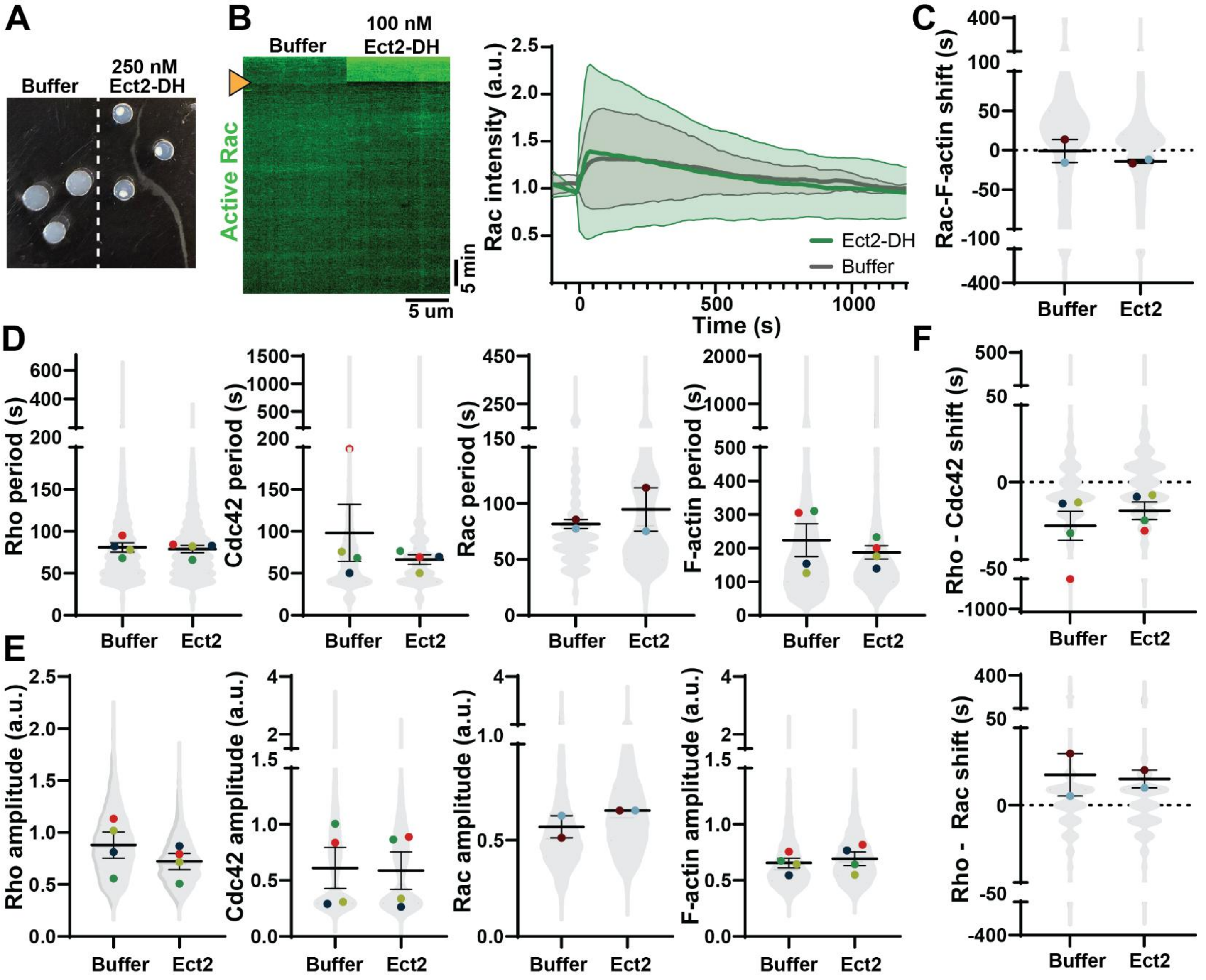


**Supplemental Figure 5. Wave dynamics following Ect2-DH treatment.**

**(A)** Gelation-contraction assay of interphase egg extract. Addition of Ect2-DH to interphase egg extract hyperactivates contractility, as seen by the concentration of light scattering particles in the center of the droplet. **(B)** *Left:* Representative kymograph of Rac intensity after 100nm Ect2-DH or vehicle control treatment. Arrow indicates time of addition. *Right:* Quantification of mean Rac intensity over time. Solid line represents mean, shaded area represents SE. n = 2 movies, 2 extract preps from 2 frogs, and 4 ROIs. Time = 0 indicates time of addition. **(C-E)** Quantification of GTPase and F-actin period, amplitude, and temporal shift following Ect2-DH or vehicle treatment. Mean ± SE is shown; violin represents all ROIs analyzed in all movies; dots indicate means of individual experiments. Dot color indicates replicates. Rho: n = 8 movies, 4 extract preps from 4 frogs, 6306 ROIs. Cdc42: n = 8 movies, 4 extract preps from 4 frogs, 6306 ROIs. Rac: n = 4 movies, 2 extract preps from 2 frogs, 3036 ROIs.

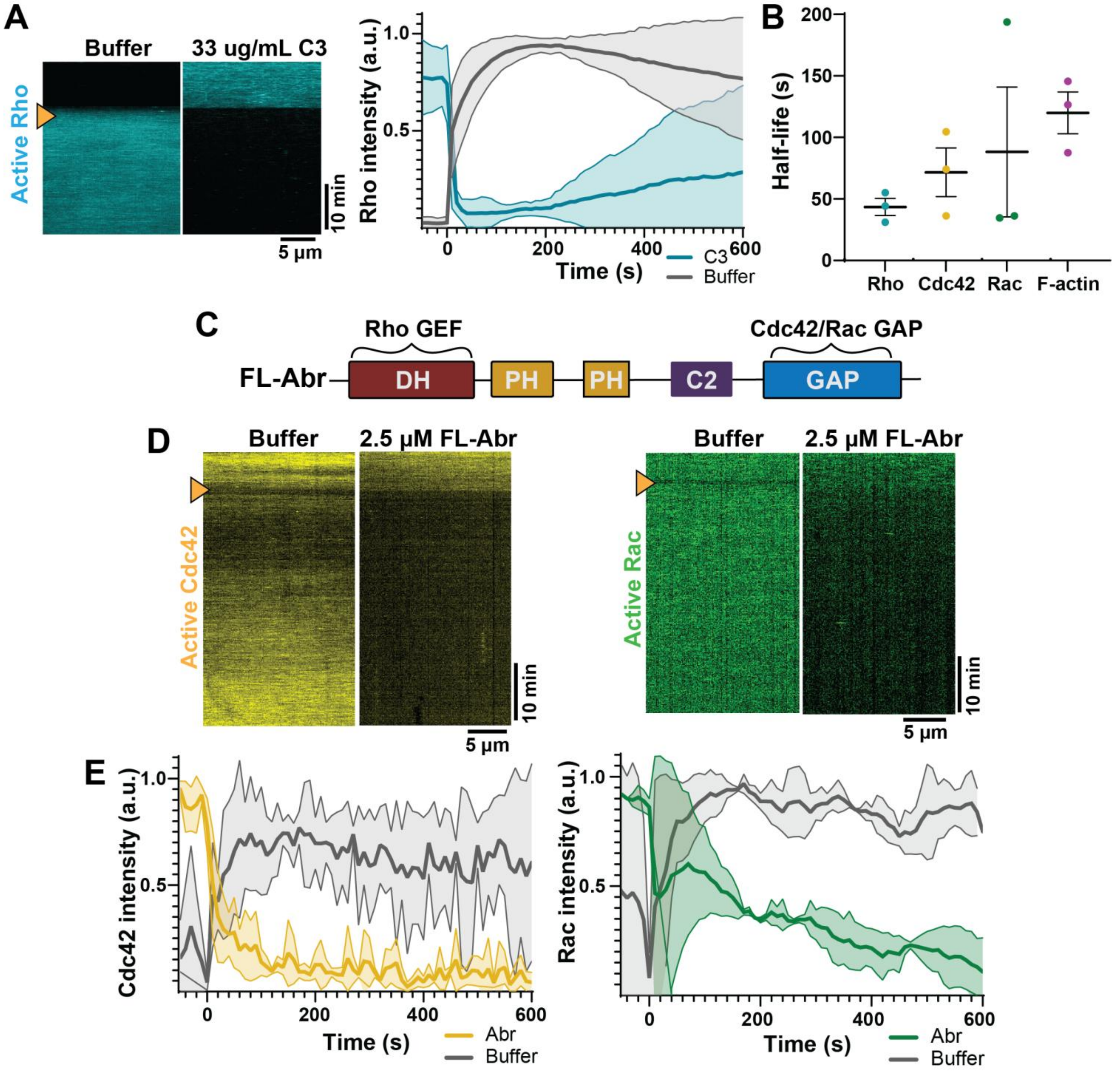


**Supplemental Figure 6. GTPase inhibition by C3 or Abr.**

**(A)** *Left:* Representative kymographs of Rho following 33 µg/mL C3 (Rho inhibitor) or vehicle control treatment. Arrow indicates time of addition. *Right:* Rho intensity over time following C3 addition (t = 0). n = 3 movies, 3 extract preps from 3 frogs, 6 ROIs. Solid line represents mean, shaded area represents SE. **(B)** Half lives of GTPases and F-actin intensity following C3 treatment. Mean ± SE is shown; dots are means of individual experiments. **(C)** Domain map of full length (FL) Cdc42/Rac inhibitor, Abr. **(D)** Representative kymographs of Cdc42 or Rac following 2.5 µM Abr or vehicle control treatment. Arrow indicates time of addition. **(E)** Cdc42/Rac intensity over time following Abr addition (t = 0). Solid line represents mean, shaded area represents SE. n = 3 movies, 3 extract preps from 3 frogs, and 6 ROIs.

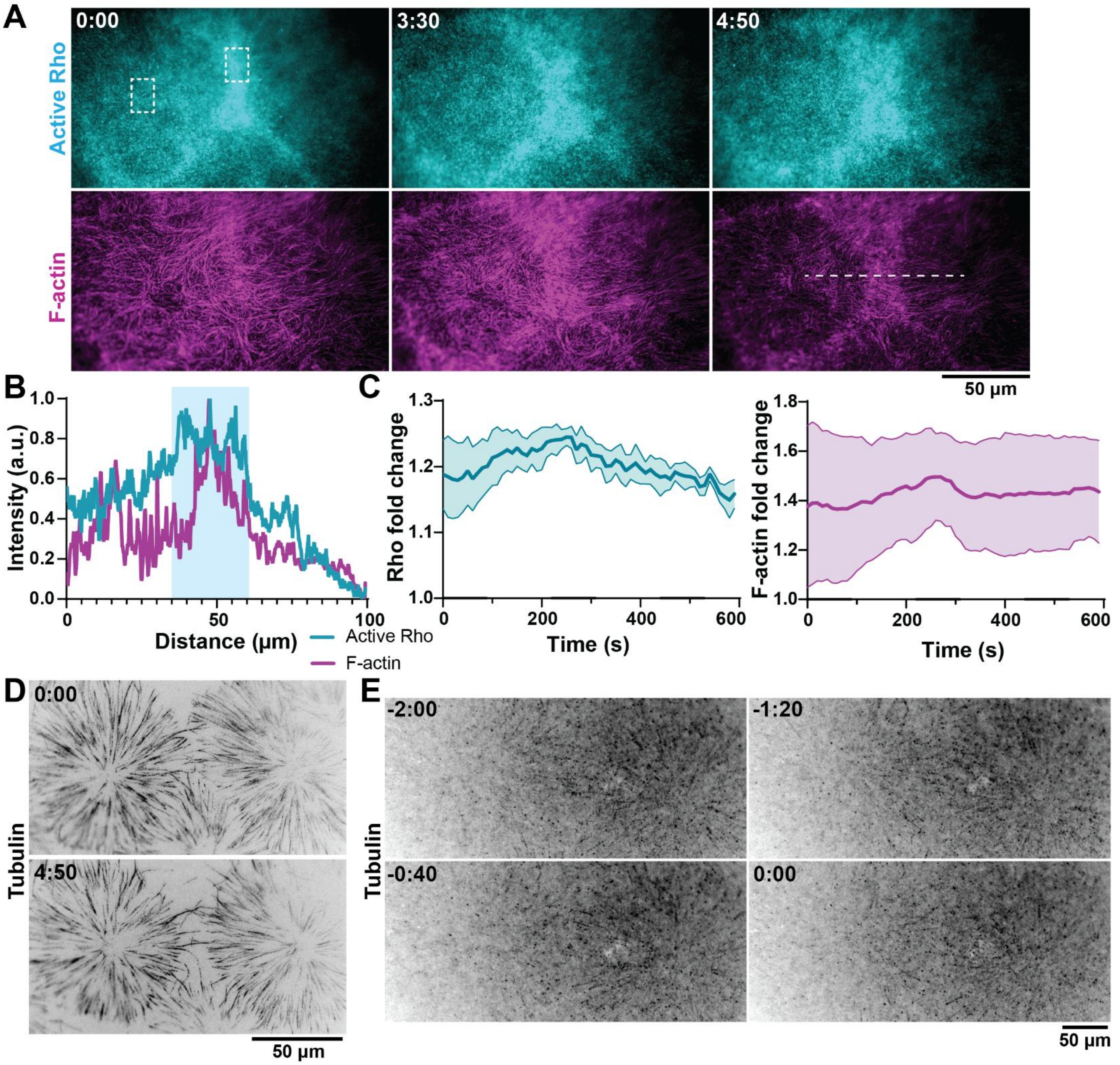


**Supplemental Figure 7. F-actin is enriched at microtubule plus-end overlaps.**

**(A)** Representative micrographs of Rho and F-actin enrichment between microtubule asters. Dotted line indicates region used for intensity profile shown in **(B)**. Boxes indicate example regions used for fold change calculations in **(C).** Time is shown in min:sec. **(B)** Intensity profile Rho and F-actin in the presence of microtubule overlap zones. Blue box indicates "division plane". **(C)** Quantification of fold change of Rho (left) or F-actin (right) at the "division plane" relative to the "non-division plane" over time. Fold change > 1 indicates increased activity at "division plane"; fold change < 1 indicates decreased activity at "division plane". Solid line represents mean, shaded area represents SE. n = 3 extract preps from 3 frogs, and 6 ROIs. **(D)** Micrographs of tubulin for experiment shown in **(A).** First and last timepoints are shown. **(E)** Micrographs of tubulin for experiment shown in Figure 6C. Time is in min:sec and corresponds to timepoints shown in Figure 6C.
